# Condensin reorganizes centromeric chromatin during mitotic entry into a bipartite structure stabilized by cohesin

**DOI:** 10.1101/2022.08.01.502248

**Authors:** Carlos Sacristan, Kumiko Samejima, Lorena Andrade Ruiz, Maaike L.A. Lambers, Adam Buckle, Chris A. Brackley, Daniel Robertson, Tetsuya Hori, Shaun Webb, Tatsuo Fukagawa, Nick Gilbert, Davide Marenduzzo, William C. Earnshaw, Geert J.P.L. Kops

## Abstract

The Structural Maintenance of Chromosomes (SMC) complexes cohesin and condensin establish the 3D organization of mitotic chromosomes^1–3^. Cohesin is essential to maintain sister chromatid pairing until anaphase onset^4^, while condensin is important for mitotic centromere structure and elastic resistance to spindle forces^5–8^. Both complexes are also important to form productive kinetochore-spindle attachments^6, 8, 9^. How condensin and cohesin work together to shape the mitotic centromere to ensure faithful chromosome segregation remains unclear. Here we show by super-resolution imaging, Capture-C analysis and polymer modeling that vertebrate centromeres are partitioned into two distinct condensin-dependent subdomains during mitosis. This bipartite sub-structure is found in human, mouse and chicken centromeres and also in human neocentromeres devoid of satellite repeats, and is therefore a fundamental feature of vertebrate centromere identity. Super-resolution imaging reveals that bipartite centromeres assemble bipartite kinetochores with each subdomain capable of binding a distinct microtubule bundle. Cohesin helps to link the centromere subdomains, limiting their separation in response to mitotic spindle forces. In its absence, separated bipartite kinetochores frequently engage in merotelic spindle attachments. Consistently, uncoupling of centromere subdomains is a common feature of lagging chromosomes in cancer cells. The two-domain structure of vertebrate regional centromeres described here incorporates architectural roles for both condensin and cohesin and may have implications for avoiding chromosomal instability in cancer cells.

---

During cell division, spindle microtubules attach to chromosomes at centromeres, specialized regions responsible for assembling the kinetochore and specified by the presence of the histone H3 variant CENP-A^10, 11^. Vertebrate chromosomes have ‘regional’ centromeres, which are variably sized and are usually enriched for satellite repeats, consisting of high-copy number tandemly repeated DNA sequences^12^. Satellite repeats, however, are not essential for centromere formation, as evolutionarily new centromeres occupy non-repetitive regions in several species, such as chicken^13^, and neocentromeres are observed in humans in rare cases^12^. The core centromere, defined as the region enriched in CENP-A chromatin containing CENP-C, is flanked by pericentromeres devoid of CENP-A. CENP-A nucleosomes in the core centromere are relatively scarce^14^ and are interspersed between canonical Histone H3-containing nucleosomes^15^. Folding of centromeric chromatin involves the packing of CENP-A nucleosomes into a yet unknown higher order organization^11, 16^. Electron microscopy analysis suggests that CENP-A packing is remarkably complex and probably involves different levels of suborganization^17^. Current models propose that centromeric chromatin may be folded in loops^18^, solenoids^15^ or as a layered sinusoid^19^. Centromere compaction is essential to preserve kinetochore integrity, allow biorientation and guarantee faithful chromosome segregation.

Centromeres remain as compact regions of heterochromatin associated with the 16 subunit CCAN complex throughout interphase^10, 11, 16^. They compact further during mitotic entry as they become enriched in cohesin and condensin, two ring-shaped complexes belonging to the Structural Maintenance of Chromosomes (SMC) family of proteins. SMC complexes are the main drivers of the 3D chromatin organization^1, 2^ and important regulators of centromeres. Centromeric cohesin keeps sister chromatids tethered during cell division^4, 20^, while condensin provides stiffness to the (peri)centromere^5, 7^. Both are required for kinetochore integrity and to prevent the formation of merotelic attachments^6, 8, 21, 22^. Despite the importance of SMC complexes in preserving the integrity of the centromere, how condensin and cohesin shape centromeric chromatin remains a fundamental question of chromosome biology.

## Regional centromeres in vertebrates are organized in two main subdomains

To better understand the structural organization of the core centromere, we performed super resolution (expansion) microscopy (ExM)^23^ of RPE-1 cells immunostained for CENP-A. Consistent with previous multi-unit centromere architecture models^15, 18, 24^, we observed that the core centromere is subdivided into discrete linked domains (Fig. 1a-c and S1a-d). Quantification of the number of subdomains per centromere revealed that over 50% of the centromeres exhibited a bipartite organization (Fig 1d and S1a), with an average inter-subdomain distance (centre-to-centre) of ∼150 nm (∼680 nm in 4.5-fold expanded samples - Fig 1e). Importantly, a similar organization of CENP-A was seen by STED microscopy on fixed samples and by iSIM microscopy on living cells expressing mCherry-CENP-A (Fig. S1e-h), ruling out artifacts from gel expansion, fixation conditions or antibody staining. When analysing preparations of stretched chromosomes, we noted that centromeres tended to split into two subdomains that each associated with a proximal chromosome arm (Fig. 1f), similar to previous observations for the first description of the CENP antigens^25^. ExM imaging of normal chromosomes likewise showed, in cases where the shape of the chromosome was clearly visible, that the continuity of the CENP-A region was interrupted by the primary constriction (inset in Fig. 1a).

**Fig. 1.**
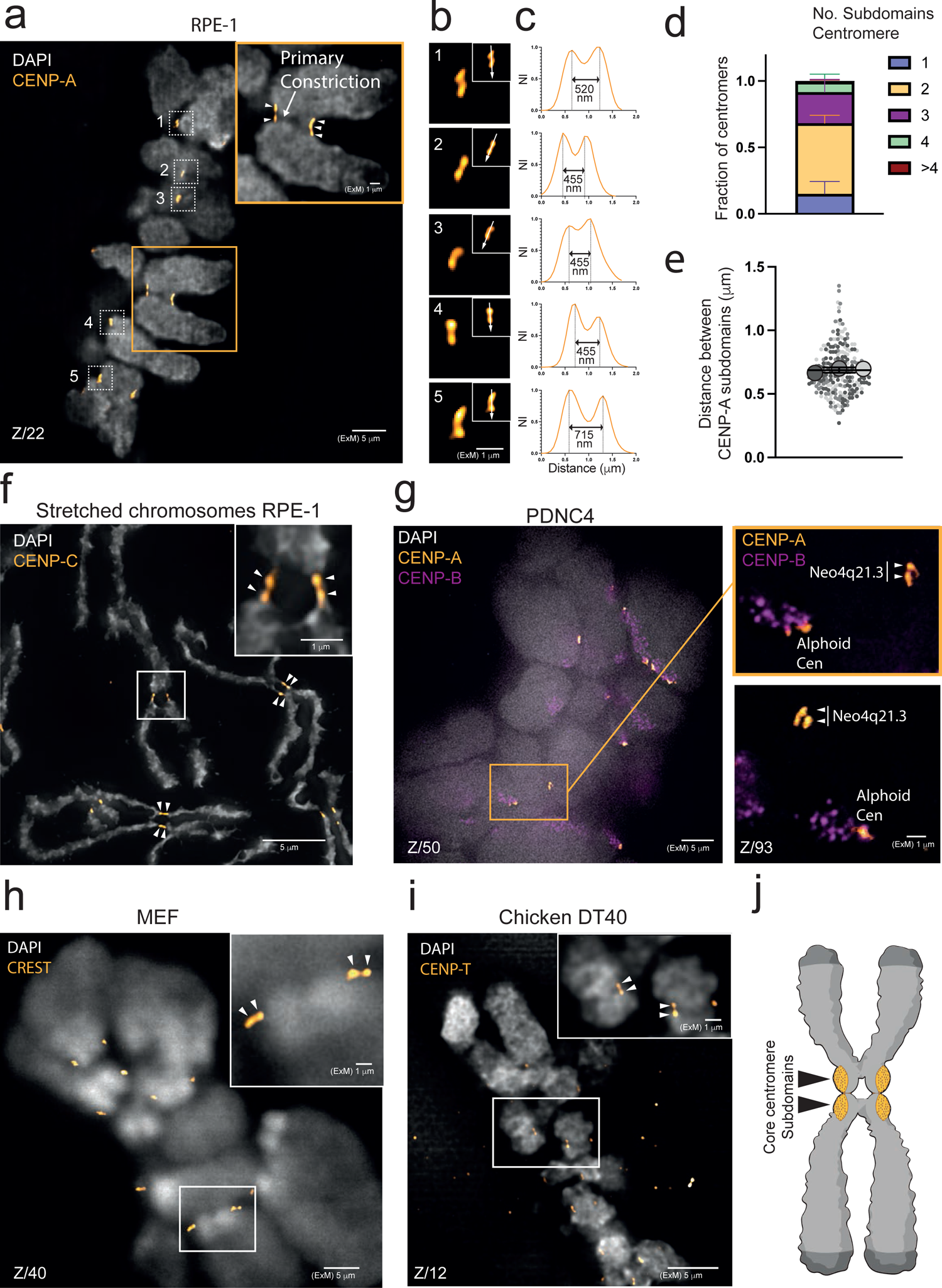
Subunit organization of the regional centromere of vertebrates. **(a-c)** a) Representative ExM image of CENP-A in RPE-1 cells in metaphase. Arrowheads in the inset (orange box) indicate CENP-A subdomains. b) Magnifications of the centromeres enclosed in the dotted boxes in (a); c) Line intensity profiles across centromere subdomains (arrows shown in b). The distance between peaks is indicated. NI= Normalised Intensity. **(d)** Quantification of the fraction of centromeres per cell that show the indicated number of CENP-A subdomains in RPE-1 cells (mean ± s.d of 4 independent experiments (*n*=35 cells). **(e)** Quantifications of the distance between CENP-A subdomains in bipartite centromeres in ExM images of RPE-1 cells (mean ± s.d. of 3 independent experiments. Each big dot represents the mean of an independent experiment. Each small dot represents a single centromere: *n*=240 centromeres from 35 cells). **(f)** Immunostaining of CENP-C in preparations of stretched chromosomes. Magnification of the boxed area is shown in the inset. Arrowheads indicate CENP-C subdomains. **(g)** Representative ExM image of CENP-A and CENP-B in PDNC4 cells harboring a neocentromere in chromosome 4 (Neo4q21.3)^26^. Magnifications of the orange box and a second region containing the sister neocentromere are shown on the right. Neo4q21.3 is recognised by the lack of CENP-B signal and canonical centromeres containing a-satellite repeats are indicated (Alphoid cen). Arrowheads indicate CENP-A subdomains in Neo4q21.3. **(h)** Representative ExM image of a mouse embryonic fibroblast (MEF) immunostained with a CREST antibody. Magnification of the boxed area is shown in the inset. Arrowheads indicate CREST subdomains. More examples from the same cell are shown in Fig. S1i. **(i)** ExM image of a chicken DT40 B cell immunostained with a CENP-T antibody. Magnification of the boxed area is shown in the inset. Arrowheads indicate CENP-T subdomains. **(j)** Cartoon depicting the organization of the core centromere, with two subdomains (represented in orange), each associated to one chromosome arm. In all images, Z specifies the plane of the Z-stack.

Importantly, the bipartite organization of centromeres was not limited to satellite repeat-containing centromeres in human cells. ExM imaging of a patient-derived neocentromere that formed on non-repetitive sequences of chromosome 4 (PDNC4)^26^, of mitotic mouse embryonic fibroblasts (with telocentric chromosomes)^27^, and of chicken DT40 B cells (which have both repetitive and non-repetitive centromere sequences)^13^ also showed bipartite centromere organisation (Fig. 1g-i and S1i). Thus, centromeric chromatin has a conserved, bipartite higher-order organisation with two major domains associated with proximal chromosome arms regardless of underlying sequence and chromosomal position (Fig. 1j).

## Bipartite kinetochore subdomains independently bind microtubule bundles

To examine how the subunit organization of the centromere impacts kinetochore function, we performed ExM on RPE-1 cells immunostained for α-tubulin and the inner kinetochore component CENP-C. Although kinetochore morphology generally looked more complex and fragmented than the CENP-A-associated core centromere (e.g. Example 1 in Fig 2a,b and Example 4 in Fig. S2a,b), the underlying bipartite configuration of the centromere was evident in many kinetochores (Fig 2a,b, S2a-c). Strikingly, close inspection of kinetochore-microtubule interactions revealed that kinetochore fibers (k-fibers) rarely appeared as one compact bundle but instead were often comprised of distinct sub-bundles that each associated with an individual kinetochore subdomain (Fig 2a,b and S2a-c). This is consistent with previous observations of *Heamanthus* (african blood lily) cell mitosis^28, 29^.

**Fig. 2.**
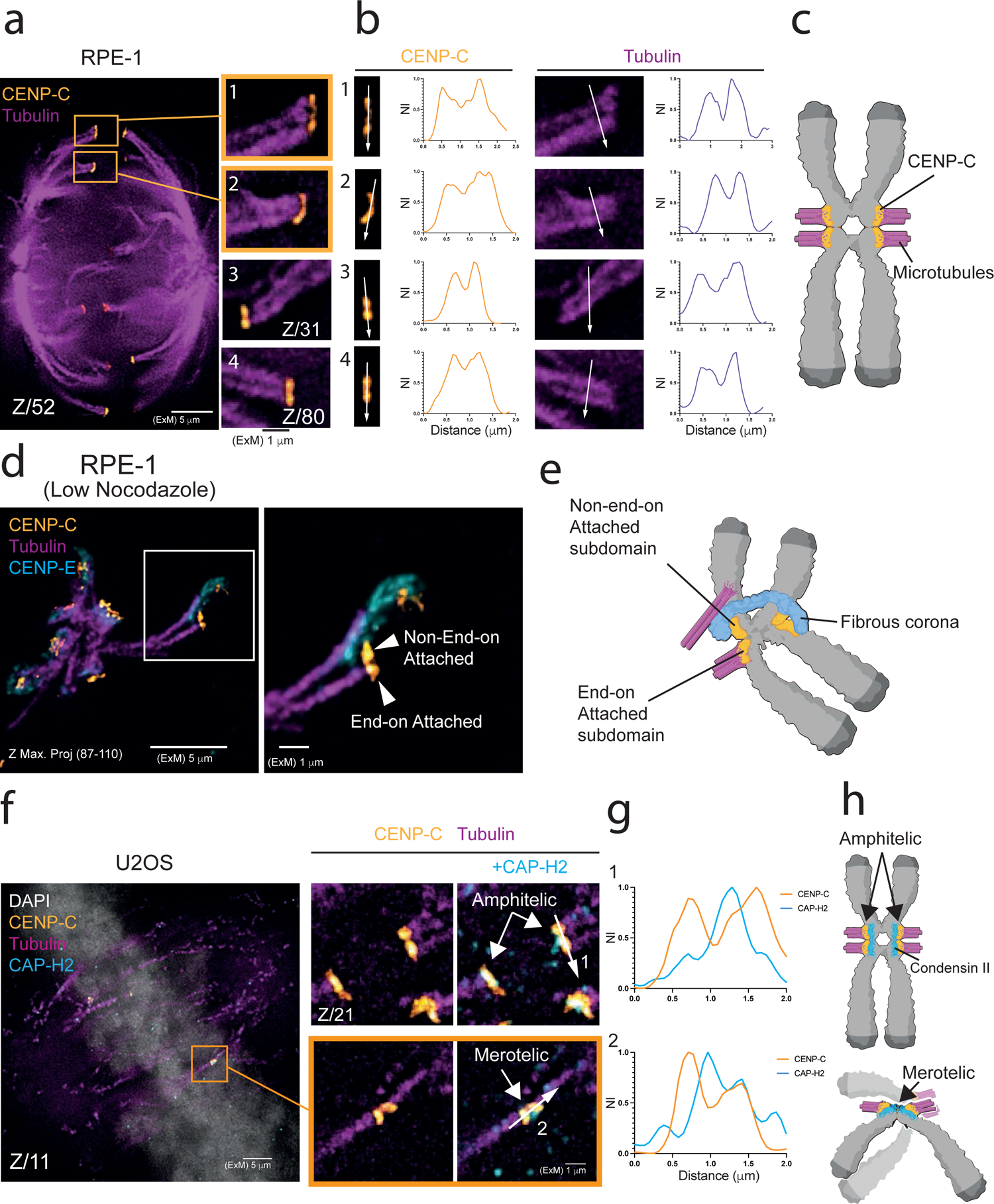
Kinetochore subdomains bind independent microtubule bundles. **(a-c)** a) Representative ExM image of α-tubulin and CENP-C in RPE-1 cells in metaphase. Magnifications of the orange boxes and other regions from the same cell are shown on the right. Cells were cold-treated before fixation. More examples from the same cell are shown in Fig. S2a; b) Line intensity profiles across kinetochore subdomains (arrows in CENP-C) and k-fibers (arrows in tubulin). NI= Normalised Intensity. c) Cartoon depicting the ability of kinetochore subdomains to bind independent bundles of microtubules. **(d,e)** d) ExM image of an aster of microtubules formed in an RPE-1 cell treated with a low concentration of nocodazole (330 nm), and kinetochores immunostained with CENP-C and CENP-E antibodies. The magnification shows a bipartite kinetochore, where one of the subdomains is end-on attached and the other is laterally attached through the fibrous corona of the kinetochore (marked by CENP-E^30^). Image is a maximum intensity projection of the indicated planes; e) Cartoon depicting the kinetochore shown in (d). **(f-h)** f) ExM image of a U2OS cell immunostained with the indicated antibodies. Magnifications show the merotelic attachment boxed in the main image and an amphitelic attachment found in the indicated plane of the same cell. g) Line intensity profiles across centromere subdomains (arrows shown in f). h) Cartoon depicting the two situations shown in (f). NI= Normalised Intensity. In all images, Z specifies the plane of the Z-stack.

These observations suggest that kinetochore subdomains can work autonomously to bind microtubules. Supporting this, in cells compromised for spindle assembly by treatment with low dose (330 nM) nocodazole, we observed kinetochores in which one subdomain was end-on attached while the other remained covered by the fibrous corona^30^, an outer kinetochore structure that is removed when end-on attachments are formed^31^ (Fig. 2d,e). Bipartite kinetochores were not a result of fragmentation caused by microtubule forces, since they were apparent also in cells that had entered mitosis in the presence of high dose (3.3 μM) nocodazole (Fig. S2d).

We next wondered if bipartite kinetochores can form merotelic attachments, an erroneous attachment configuration in which a single kinetochore engages microtubules from both spindle poles, often resulting in lagging chromosomes in anaphase^21^. In high-resolution live-cell imaging of anaphase U2OS cells, an osteosarcoma cell line with high rates of lagging chromosomes^32^, we observed that around 50% of lagging chromatids showed split CENP-A signals (Fig. S3a-d and Supplementary Movie 1). ExM imaging further revealed merotelic attachments in which the two subdomains of a single kinetochore attached to opposite sides of the spindle (Fig. 2f,g). Immunostaining for the condensin II subunit CAP-H2 verified that these represented subdomains of one kinetochore rather than two sister kinetochores: CAP-H2 is normally found close to individual core centromeres^8^ (Fig. 2f,g), and in the merotelic chromatids could be seen between kinetochore subdomains (Fig. 2f,g). We conclude that centromere subdomains work as independent microtubule-binding units and can potentially form merotelic attachments (Fig. 2h).

## Bipartite organisation of the chicken Z-centromere revealed by Capture-C analysis

Chicken DT40 cells have a single Z chromosome with a non-repetitive centromere (Zcen)^13^. In this system, we previously observed a substantial change of chromatin higher-order packing at centromeres during a highly synchronous transition from G_2_ into mitosis in relatively low resolution Hi-C data^3^ (Fig. 3a and S4a). To better understand this change, we constructed a high-resolution interaction map of the Zcen in G_2_ and mitosis using Capture-C analysis. Capture-C measures DNA proximity from multiple locations of interest (viewpoints) in cells and can resolve structural details over genomic regions the size of chicken centromeres^33^ (Fig. 3b). To analyse the Capture-C data, we assessed the directionality of interactions over a range of distances (e.g. 3-10 kb to 3-1000 kb) from each viewpoint.

**Fig. 3.**
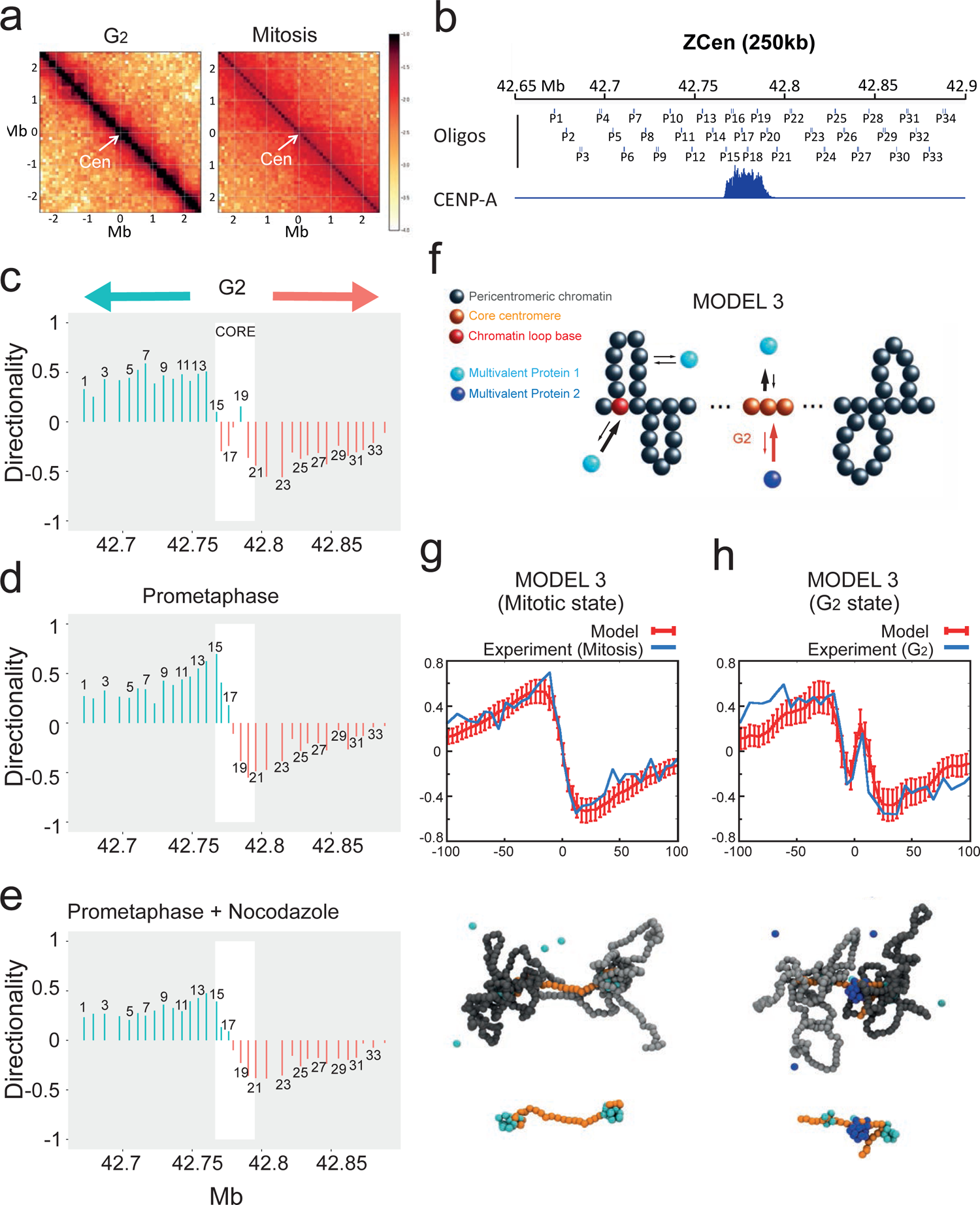
Bipartite mitotic centromeric chromatin organisation of Zcen. **(a)** Hi-C map of 2.5 MB region surrounding Zcen (arrow) in G_2_ or in late prometaphase (T=30 min after release from 1NMPP1). 100 kb resolution. Data is taken from (^3^). **(b)** Positions of Capture oligos (P1-P34) surrounding Zcen and the CENP-A ChIP-seq data to indicate the region of Z core centromere of WT CDK1^as^ subclone cell line used for this study. **(c-e)** Directionality of interactions at each view point in wild type G_2_ cells (c), wild type late prometaphase cells (d), wild type late prometaphase cells treated with nocodazole (e). Core centromeric region (defined by the presence of CENP-A) is marked by white box. Asymmetry in interaction is depicted by green upwards bar (more interactions towards p arm) and by orange downwards bar (more interactions towards q arm). X axis shows genomic DNA position in Z chromosome. Value on the Y axis is the natural log of the number of interactions towards the p arm divided by the number of interactions towards the q arm; only interactions with positions within a distance of 3-250 kbp of the viewpoint are included. **(f)** Polymer physics model 3 of the centromere, considering two types of multivalent chromatin-binding proteins (MP1 and MP2) (or bridges, representing SMC proteins^34^). MP1 binds pericentromeric and centromeric chromatin, with a higher affinity for the former (represented by the length and thickness of the arrows). MP2 binds the core centromere and is present only in G_2_. This model also includes chromatin loops formed in the pericentromeric arms (e.g. SMC-mediated by loop extrusion^3,36^), which create a bottlebrush topology. Note that the affinity is highest at the base of chromatin loops, an example of which is coloured in red. **(g,h)** Capture-C-like asymmetry plots (upper panel) and typical 3D configurations obtained in equilibrium (two lower panels) in a mitotic- (g) and G_2_-like (h) state of model 3. In the 3D models, dark and light gray beads denote the two pericentromeric arms, orange beads denote the core centromere, and cyan and blue beads denote MP1 and MP2, respectively. The mitotic-like state has typically a bipartite structure with MP1 clusters separated by a largely unstructured core centromere (g). In the G_2_-like state, MP2 forms an additional cluster at the core centromere (h). The transition between the G_2_ and the mitotic states can be triggered, for instance, by decreasing the affinity between the MP2 and the core centromere (See methods and figure S4b for more details).

Capture-C analysis of a region of ∼250 kb flanking the Zcen revealed that while core centromeric chromatin in G_2_ cells tended to interact locally within the core CENP-A-enriched domain, the flanking pericentromeric chromatin preferentially interacted away from the core centromere (Fig. 3c). The core centromere region thus forms a boundary limiting interactions between p- and q-arm pericentromeres. Capture-C analysis further revealed a striking change in chromatin folding at the Zcen as G_2_ cells entered mitosis (Fig. 3d). In contrast to an apparent internal looping structure in G_2_ (Fig. 3c), mitotic centromere chromatin adopted a striking left-right bipartite asymmetry centered within the core (Fig. 3d). This did not require kinetochore-microtubule interactions, as nocodazole-treated cells showed a similar organisation (Fig. 3e). Reproducibly, the directionality of left-right interactions was highest at the boundary between the CENP-A core and pericentromere and decreased for viewpoints moving away from the core (Fig. 3d).

In summary, the core centromere is reorganized as G_2_ cells enter mitosis. This involves the loss of strictly intra-core interactions and the establishment of a sharp boundary that prevents interactions between chromatin domains on either side of the boundary and favours asymmetrical contacts outwards towards the adjacent pericentromere regions. Thus, core centromeric chromatin adopts a bipartite configuration upon mitotic entry, consistent with our microscopy observations.

## Polymer modelling of SMC activities in the (peri)centromere

To better understand chromatin folding we used polymer modelling to investigate potential mechanisms that could lead to a bipartite 3D organisation of pericentromeric chromatin. We considered three key ingredients (see methods and figure legends for details, and Fig. 3f): i) multivalent proteins able to bind either pericentromeric chromatin or centromeric chromatin (bridges, for instance provided by SMC complexes^34^); ii) a nonequilibrium biochemical reaction of the proteins (e.g. ATP hydrolysis), enabling them to switch from a binding to a non-binding state and vice-versa^35^; iii) (stable) loops at random positions in the two pericentromere regions, modelling condensin- or cohesin-mediated loops^3, 36^.

This analysis successfully reproduced the Capture-C profile seen in G_2_ as well as the dramatic change that occurs on mitotic entry. When including bridges alone, the model approximated the overall shape of the Capture-C directionality plot, provided that the binding affinity of proteins to pericentromeres was larger than to the core centromere (Model 1, Fig. S4b). In this case, but not when binding affinities were swapped (Model 0, Fig. S4b), chromatin formed a spherical globule with the two pericentromere regions folded into two different hemispheres and the centromere arranged on the surface. Increasing the rate of switching led to the splitting of the globular structure in a significant portion of structures in the population, whilst retaining a significant asymmetry in the contact pattern as observed by Capture-C (Model 2, Fig. S4b,c). Subsequent introduction of pericentromere loops (e.g. cohesin or condensin-mediated^37^) resulted in a more robust bipartite 3D structure, with enhanced directionality in the contacts and sharper transition between the two sides of the centromere – closely fitting our Capture-C data (Model 3, Fig. 3f,g and S4c). Interestingly, the addition in model 3 of a second bridging activity with preferential binding to the core centromere (e.g. centromeric cohesin) resulted in three main chromatin clusters, accurately reproducing the more complex directionalities observed in G_2_ (Fig. 3f,h and S4c).

## (peri)centromeric distribution of SMC complexes

Modeling most closely recapitulated our Capture-C and microscopy observations, when it incorporated a mechanism for bipartite organisation of pericentromeres flanking the core centromere that depended on (i) SMC-mediated chromatin bridging and (ii) SMC-mediated chromatin looping flanking the centromere core. This inspired us to assess the (peri)centromeric distribution of condensin and cohesin complexes. We therefore generated a SMC3-TurboID^38^ knock-in in the background of a SMC2-mAID-mCherry^39^ cell line. To visualise cohesin (SMC3) distribution, we imaged biotinylated substrates in nocodazole-treated cells using partial gel expansion (∼2x). This approach achieved enhanced spatial resolution of cohesin while retaining sufficiently high signal intensity. Although this did not resolve centromere subdomains, it did reveal cohesin distribution in two distinct (peri)centromeric locations (Fig. 4a). A main pool of cohesin accumulated between sister (peri)centromeres and frequently appeared as two foci flanking the kinetochore along the chromosome axis, marked by SMC2 (asterisks in Fig. 4a,b). This distribution is reminiscent of that of the inner centromere protein INCENP^40^, and is consistent with the expected role of cohesin in tethering sister chromatids near pericentromeric regions. A second pool of cohesin was observed proximal to the CENP-A region and partly overlapped with condensin (yellow arrows in Fig. 4a,b), which accumulates to higher levels at the core and/or pericentromeric region^8, 41^ (Fig. S5a,b).

**Fig. 4.**
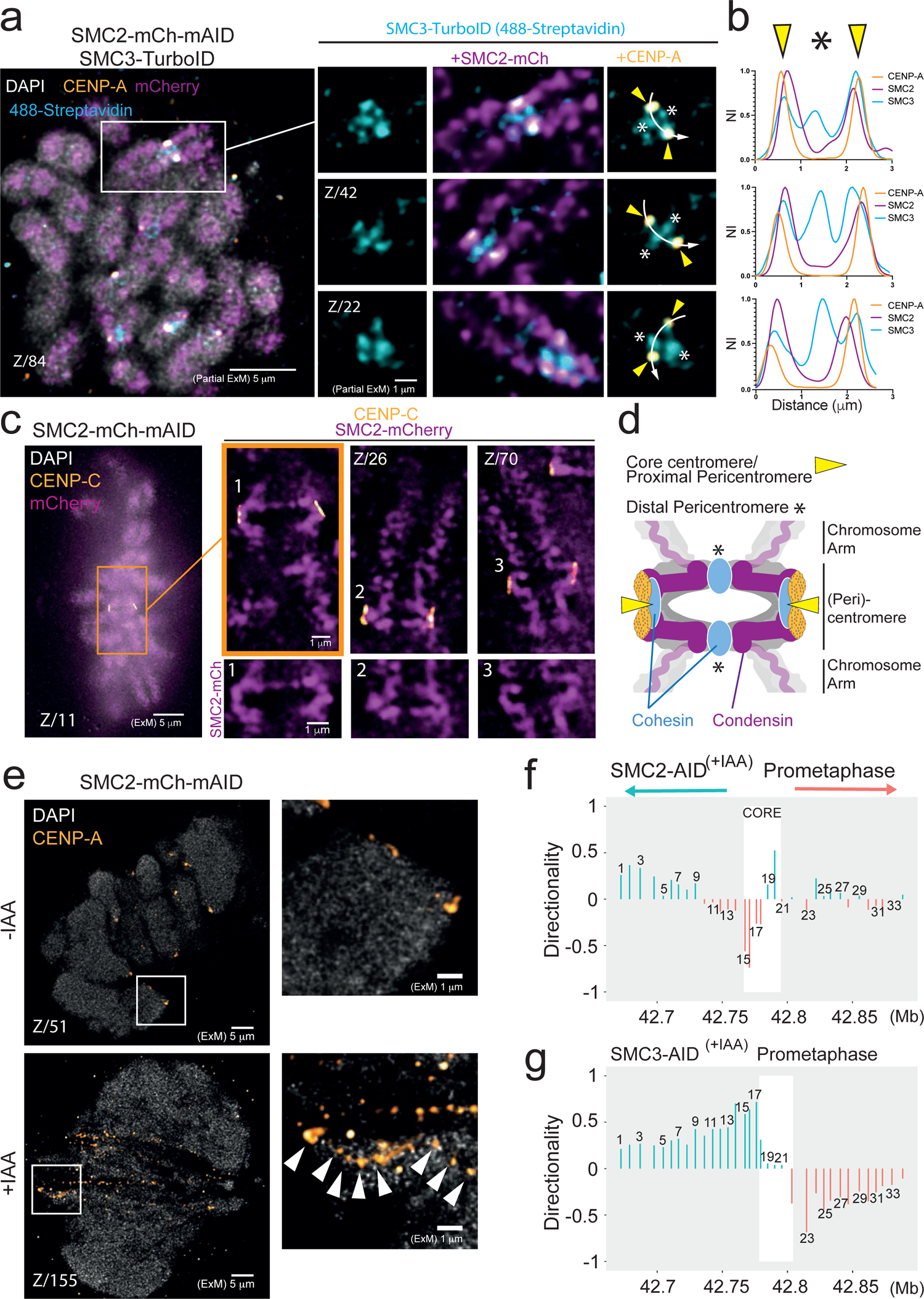
Condensin drives the partitioning of the centromere in mitosis. **(a,b)** (a) Partially expanded image of an HCT-116 cell expressing SMC2^-^mAID-mCherry and SMC3-TurboID, and stained as indicated. Cells were synchronized with thymidine and released in nocodazole. Biotin was added to the media 30 min before fixation. Biotinylated substrates were visualized with Streptavidin-AF488. Magnifications of the orange box and other regions from the same cell are shown on the right. (b) Line intensity profiles across sister centromeres (arrows in b). Yellow arrowheads indicate the pool of cohesin situated proximal to the core centromere and asterisks indicate the pool of cohesin situated at the inner centromere. NI= Normalised Intensity. **(c,d)** c) ExM image an HCT-116 cell expressing SMC2^-^mAID-mCherry and immunostained with the indicated antibodies. Magnifications of the orange box and other regions from the same cell are shown on the right. The lower insets show zooms of the indicated centromeres. Note the C-shape of the (peri)centromere and a bridge of SMC2 connecting both sister (peri)centromeres in zoom 1. **(d)** Model integrating the observed distributions of SMC complexes in the (peri)centromere. **(e)** ExM images of CENP-A in HCT-116^SMC2-mAID-mCherry^ cells prepared in the absence or presence of auxin (IAA). Magnifications of the boxed areas are shown on the right. Arrowheads indicate highly fragmented CENP-A subdomains. **(f,g)** Directionality of interactions at each view point in SMC2-AID (f) and SMC3-AID (g) late prometaphase cells treated with auxin. Asymmetry in interaction is depicted by green upwards bar (more interactions towards p arm) and by orange downwards bar (more interactions towards q arm). X axis shows genomic DNA position in Z chromosome. Value on the Y axis is the natural log of the number of interactions towards the p arm divided by the number of interactions towards the q arm; only interactions with positions within a distance of 3-250 kbp of the viewpoint are included. Note: the core centromeric region is marked by a white box. In SMC3-AID cells this is shifted towards the q arm by ∼10 kb compared to wt and SMC2-AID cells due to centromere migration in this clone.

In fully expanded samples of metaphase cells, (peri)centromeric SMC2 (in these images defined as the chromosomal regions flanking the kinetochore) formed a C-shape (Fig. 4c, zooms). This results from the tension created across the pericentromere in the attached state, as (peri)centromeric SMC2 appeared compacted in nocodazole-treated cells (Fig. 4a). The edges of the “C” tended to converge towards those of the sister chromatid (Fig. 4c, zooms). In some cases, sister pericentromeres appeared connected (Fig. 4c, zoom 1). Based on the cohesin distribution (asterisks in Fig. 4a,b), we speculate that the convergence of sister pericentromeres results from the establishment of sister chromatid cohesion at the distant pericentromere^42^ (Fig. 4d). Together, these analyses indicate that condensin and two pools of cohesin occupy distinct domains in (peri)centromeric chromatin (Fig. 4d).

## Condensin drives the partitioning of the centromere in mitosis

Condensin is required to provide compliance to centromeric heterochromatin in response to microtubule forces^5–8, 43^. Consistent with this, CENP-A appeared stretched and heavily fragmented in metaphase HCT-116^SMC^^2-mAID-mCherry^ cells depleted of the condensin subunit SMC2 (Fig. 4e). Capture-C analysis of DT40 cells acutely depleted of condensin showed that while G_2_ cells maintained a wild type interaction profile (Fig. S5c,d), the bipartite structural transformation in centromeric chromatin during mitotic entry was abolished (Fig. 4f). This was most apparent in cells in which microtubules were absent (Fig. S5e). In those cells, the mitotic Zcen had an organization indistinguishable from that of G_2_ cells (Fig. S5d). In contrast to condensin depletion, lack of cohesin (SMC3 depletion) did not alter the folding of the Zcen in G_2_ or mitosis (Fig. 4g and S4c,f-h). Thus, condensin but not cohesin is essential for establishing the bipartite centromere organization in mitotic cells.

## Cohesin stabilises centromere subdomains and promotes amphitelic spindle attachments

To build on the predictions from polymer modeling and our observations of a substantial amount of cohesin at or near the core centromere, we examined a potential role for cohesin in maintaining the architectural integrity of the centromere^42, 44^. Degron-mediated depletion of Sororin^45^, a protector of centromeric cohesin from Wapl-mediated release^46, 47^, produced a pronounced cohesion fatigue phenotype (Supplementary Movie 2), with single chromatids misaligned at spindle poles (arrowheads in +IAA condition in Fig. 5a). Some chromatids, however, remained congressed (rectangle in +IAA condition in Fig. 5a), but their centromeres were often split in two (Fig. 5b,c). Mitotic entry in the presence of nocodazole prevented centromere splitting, suggesting a crucial role for spindle forces (Fig. 5c and S6a,b). Strikingly, ExM imaging of kinetochores and microtubule connections of the congressed single chromatids in Sororin-depleted cells revealed that kinetochores had often split in two resulting in single chromatid biorientation (Fig 5d and S6c,d; ∼85% of cells with obvious cohesion fatigue phenotype). This represents an extreme variant of merotelic chromosome-spindle interactions (Fig. 5f). Visualisation of CAP-H2, marking the axis of chromatids and centromeres, verified that bioriented kinetochores represented single split centromeres rather than sister centromeres, by virtue of its location between CENP-C signals (Fig 5d and S6c,d). Similar results were seen upon RNAi of RAD21, the α-kleisin subunit of cohesin (Fig. 5e). Thus, cohesin stabilises the bipartite centromere organisation in mitosis and promotes correct amphitelic orientation of sister chromatids.

**Fig. 5.**
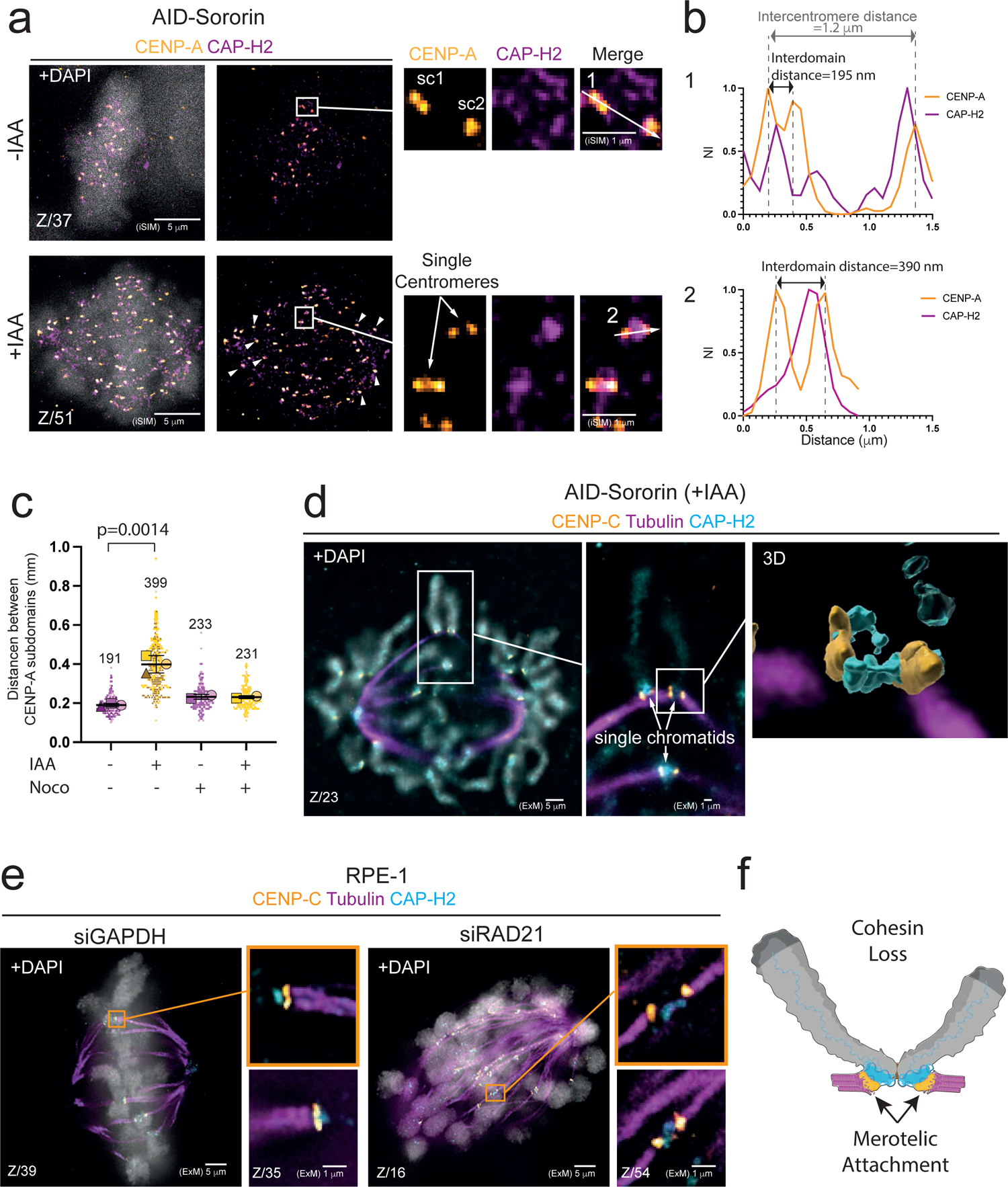
Cohesin stabilizes the bipartite centromere. **(a-c)** a) iSIM microscopy images of CENP-A and CAP-H2 in HeLa^AID-Sororin^ cells prepared in the absence or presence of auxin (IAA). Cells were released from RO-3306 for 30 minutes and treated with MG-132 for another 30 minutes before fixation. In the main image of +IAA condition, white arrowheads indicate unaligned unpaired chromatids. Magnifications of the boxed areas are shown on the right. Single centromeres (sc) are indicated; (b) Line intensity profiles of the arrows shown in a). The upper line scan crosses the two sister centromeres (arrow between sc1 and sc2), including the two subdomains of sc1. The lower line scan crosses a single centromere of an unpaired chromatid. Intercentromere and interdomain distances are indicated. NI= Normalised Intensity. c) Quantification of the distance between CENP-A subdomains in HeLa^AID-Sororin^ cells treated as indicated. Images of nocodazole-treated cells are shown in Fig. S6a. (Mean ± s.d. of 3 (-Noco) and 2 (+Noco) independent experiments. Each big dot represents the mean of an independent experiment. Each small dot represents a single centromere: -IAA-Noco: (*n*=195 centromeres from 28 cells); +IAA-Noco (*n*=301 centromeres from 33 cells); -IAA +Noco (*n*=170 centromeres from 26 cells); +IAA +Noco (*n*=164 centromeres from 26 cells). (Student’s t-test, two-tailed, unpaired. *t*=7.934, *df*=4). **(d)** ExM image of a HeLa^AID-Sororin^ cell treated with IAA and stained with the indicated antibodies. Cells were cold-treated before fixation. The central panel is a magnification of the indicated region, showing three unpaired chromatids with split kinetochores due to the formation of merotelic attachments. This is best seen in the 3D reconstruction shown in the right panel. More examples are shown in Fig. S6c,d. **(e)** ExM images of RPE1 cells treated with siGAPDH or siRAD21 and stained as indicated. Magnifications of the orange boxes and other regions from the same cell are shown on the right side of the main images. Cells were cold-treated before fixation. **(f)** Cartoon depicting the phenotype of cohesin loss, leading to the separation of kinetochore subdomains due to the formation of merotelic attachments.

## DISCUSSION

Centromeric chromatin is composed of a compact core domain enriched in CENP-A nucleosomes flanked by heterochromatic pericentromeres. Here, we find that, surprisingly, the CENP-A core domain can reproducibly be resolved into two subdomains, each of which appears to be closely associated with its adjacent pericentromere. Under some conditions further fragmentation of the CENP-A domains can be observed^18^, but a bipartite organisation is by far the most common. These CENP-A subdomains can independently bind separate microtubule bundles, and this can in some cases lead to merotelic chromosome attachments.

Our data can be explained by models in which cohesin in G_2_ cells has a significant affinity for the core centromere, and this affinity is significantly reduced when cells enter mitosis. In G_2_, cohesin is highly abundant^47^ and its binding to the core centromere can cause the region to collapse into one or more local loops (Fig. 3c). These core loops are flanked by the pericentromere, with left and right sides tending to remain separate, mainly due to entropic exclusion of loops formed by cohesin or possibly condensin II. As a result, the core centromere maintains a relatively compact heterochromatin-like organisation in interphase^48, 49^.

Upon mitotic entry, cohesin levels fall substantially^47^ and our modelling suggest that its affinity for the core centromere also decreases. Most cohesin may need to be cleared from the CENP-A core during mitosis to allow CCAN complexes to adopt a suitable folding to assemble the kinetochore plate. Next, newly activated condensin binds to the pericentromere where it is responsible for establishing the compliance (elasticity) of the heterochromatin, possibly by loop extrusion. If the CCAN complexes tend to associate with condensin or other components in the pericentromere, then treatments that tend to stretch the centromere might tend to separate the CENP-A core centromere into two domains - one proximal to each pericentromere.

A similar loop-based organization has been proposed for yeast centromeres to explain the physico-mechanical properties of the pericentromere and the observed compaction of the centromere in mitosis^44^. Interestingly, and in striking similarity to our Capture-C data, recent Hi-C analysis of yeast pericentromeres revealed a tendency of the regions flanking the core centromere to interact with the pericentromere of the same arm^42^. Since our modeling indicates that the structure of the pericentromere is a major determinant of core centromere organization, a conserved condensin-driven bottlebrush architecture of the pericentromere^44^ can therefore explain why bipartite centromeres are present along regional centromeres of a wide range of vertebrate species. Given observations of a bipartite kinetochore structure in *Haemanthus*^28, 29^, they may also be present in higher plants. If the resulting bipartite organisation for the core centromere has any function remains unclear. It might increase flexibility of this region, possibly helping to promote efficient microtubule attachment and to maintain that attachment during chromosome oscillations.

Despite its reduced levels in mitosis, cohesin is crucial for maintaining centromere integrity, linking the two centromeric subdomains and minimizing merotelic attachments. Interestingly, reduced cohesin levels have been linked to kinetochore fragmentation and merotely in ageing oocytes^22^. This function of cohesin in stabilizing the bipartite centromere may be also impaired and promote chromosomal instability in tumors harboring mutations in cohesin subunits, which are frequent in cancer^50, 51^.

In summary, our studies reveal a higher-order organization of (peri)centromeric chromatin that is formed and stabilized by condensin and cohesin. Our findings raise new fundamental questions as how other mechanisms (e.g. CCAN members^24, 52, 53^, CENP-B^54^) contribute to the suborganization of this bipartite chromatin structure, and how biorientation is regulated and achieved in the context of a bipartite kinetochore.

---

**Fig. S1. Subunit organization of the regional centromere of vertebrates. (a)** 3D reconstructions of the centromeres shown in Fig. 1a,b. **(b-d)** b) Representative ExM image of CENP-A in RPE-1 cells in metaphase; c) Magnifications of the centromeres enclosed in the boxes in b). Z indicates the specific plane shown in the magnification; d) Line intensity profiles across centromere subdomains (arrows shown in c). The distance between peaks is indicated. NI= Normalised Intensity. **(e,f)** e) Immunostaining of CENP-A in HCT-116 cells imaged by confocal and STED microscopy. Numbered squares enclose the same centromeres in both conditions. Magnifications of the boxes in the STED image are shown on the right column; f) Line intensity profiles across centromere subdomains (arrows shown in e). The distance between peaks is indicated. NI= Normalised Intensity. **(g,h)** g) iSIM live-cell imaging (lateral resolutioñ125 nm) of a U2OS cell expressing mCherry-CENP-A and H2B-mNeon. Magnifications of the orange box and other regions from the same cells are shown on the right; h) Line intensity profiles across centromere subdomains (arrows shown in g). The distance between peaks is indicated. NI= Normalised Intensity. **(i)** Representative ExM image of a mouse embryonic fibroblast (MEF) immunostained with a CREST antibody. Magnification of the boxed area is shown in the inset. Arrowheads indicate CREST subdomains. The image belongs to the same cell shown in Fig. 1h. In all images, Z specifies the plane of the Z-stack.

**Fig. S2. Kinetochore subdomains bind independent microtubule bundles. (a,b)** a) Representative ExM image of α-tubulin and CENP-C in an RPE-1 cell in metaphase. Magnifications of the numbered boxes are shown on the right. Cells were cold-treated before fixation. The image belongs to the same cell shown in Fig. 2a; b) Line intensity profiles across kinetochore subdomains (arrows in CENP-C images) and k-fibers (arrows in tubulin images). NI= Normalised Intensity. Images correspond with the kinetochores boxed in a). **(c)** Representative ExM image of α-tubulin and CENP-C in an RPE-1 cell in metaphase. Magnifications of the orange box and other regions from the same cell are shown on the right. **(d,e)** d) Representative ExM image of CENP-C and CENP-A in RPE-1 cells released from an RO-3306 arrest in the presence of a high concentration of nocodazole (3.3 μM). Magnifications of the boxed areas are shown on the right. e) Line intensity profiles across centromere subdomains (arrows in d). NI= Normalised Intensity. In all images, Z specifies the plane of the Z-stack.

**Fig. S3. Mitotic errors in U2OS cells expressing CENP-A-mCherry and H2B-mNeon. (a-b)** a) iSIM movie of a U2OS cell showing a lagging chromosome with a centromere that splits as anaphase progresses. Arrowheads indicate split CENP-A signal. See also Supplemental Movie 1. **(b-d)** Examples of U2OS cells in anaphase showing the indicated types of errors (lagging chromosomes (b) or bridges (c)), and quantification of the frequencies of each type of error (d). A total of 60 divisions were filmed in 3 independent experiments, of which 26 showed mitotic errors. Total number of observed errors=37. All images are maximum projections in Z.

**Fig. S4** Bipartite mitotic centromeric chromatin organisation of Zcen. **(a)** Hi-C map of 25 MB region surrounding Zcen (arrow) in G_2_ or in late prometaphase (T=30 min after release from 1NMPP1). 100 kb resolution. Data is taken from Gibcus *et al.*^3^ **(b)** Polymer physics models 0, 1 and 2 of the centromere. All models consider one type of multivalent protein (MP), (or bridges^34^, representing SMC proteins and other histone-binding proteins). MP binds the pericentromeric and centromeric chromatin with different affinities (represented by the length of the arrows). Capture-C-like asymmetry plots and typical 3D configurations obtained in equilibrium are shown. In the 3D models, dark and light gray beads denote the two pericentromeric arms, orange beads denote the core centromere, and cyan beads denote the MP. In model 0, MP shows a higher affinity for the core centromere. This results in an “inverted” Capture-C signal with respect to the mitotic one, and in a compact configuration that shows the core centromere buried in the structure. In model 1, MP binds more strongly to the pericentromere than to the core centromere. This leads to a qualitatively correct Capture-C signal, and to a switch in the core centromere location, which now is peripheral. In Model 2, a nonequilibrium biochemical reaction of MP is included. This allows the switching of MP and leads to a bipartite organisation in a significant portion of structures in the population, whilst retaining the asymmetry in the contact pattern as observed by Capture-C. (See methods for more details). (c) Fraction of structures in the population of models 0, 1, 2 and 3 showing a monopartite or bipartite organization.

**Fig. S5. Centromeric localization and functions of condensin and cohesin. (a,b)** Immunofluorescence image of an HCT-116 cell expressing SMC2-mAID-mCherry and stained as indicated. (b) Line intensity profiles across chromosome arms (arrows in a). NI= Normalised Intensity. **(c)** Positions of Capture oligos (P1-P34) surrounding Zcen and the CENP-A ChIP-seq data to indicate the region of Z core centromere of WT, SMC2-AID, SMC3-AID CDK1^as^ subclone cell lines used for this study. **(d-h)** Directionality of interactions at each view point in SMC2-AID G_2_ (d), SMC2-AID Late Prometaphase + nocodazole (e), and SMC3-AID G_2_ (f), treated with auxin (IAA). Also SMC3-AID G_2_ (g), and SMC2-AID late prometaphase (h) without auxin are shown as control. Asymmetry in interaction is depicted by green upwards bar (more interactions towards p arm) and by orange downwards bar (more interactions towards q arm). X axis shows genomic DNA position in Z chromosome. Value on the Y axis is the natural log of the number of interactions towards the p arm divided by the number of interactions towards the q arm; only interactions with positions within a distance of 3-250 kbp of the viewpoint are included. Note: the core centromeric region is marked by a white box. In SMC3-AID cells this is shifted towards the q arm by ∼10 kb compared to wt and SMC2-AID cells due to centromere migration in this clone.

**Fig. S6. Cohesin stabilizes the bipartite centromere. (a,b)** a) iSIM microscopy images of CENP-A and CAP-H2 in HeLa^AID-Sororin^ cells that entered mitosis in the presence of nocodazole and prepared in the absence or presence of IAA. Cells were released from RO-3306 directly in nocodazole (3.3 μM) for 1 hour before fixation. Single chromatids (sc) are indicated. Magnifications of the orange boxes and other regions from the same cell are shown on the right. Single centromeres (sc) are indicated. (b) Line intensity profiles across centromere subdomains (arrows in a). Interdomain distances are indicated. NI= Normalised Intensity. **(c,d)** (c) ExM images of HeLa^Sororin-AID^ cells prepared in the absence or presence of IAA and stained as indicated. Magnifications of the boxed areas and other regions from the same cell are shown on the right. Single centromeres (sc) are indicated; Cells were cold-treated before fixation. (d) Line intensity profiles across kinetochore subdomains or sister kinetochores (arrows in c). NI= Normalised Intensity.

**Supplementary Movie 1.** iSIM movie of an anaphase of a U2OS cell expressing H2B-mNeon (gray) and mCherry-CENP-A (orange). Note the lagging chromosome showing a centromere that splits as anaphase progresses. Stills are shown in Fig. S3a.

**Supplementary Movie 2.** Time lapse of Hela^AID-Sororin^ cells expressing H2B-mNeon and treated as indicated.

## MATERIALS AND METHODS

### Cell culture and generation of stable cell lines

RPE-1 cells were cultured in DMEM/F-12 (Gibco) supplemented with 10% Tet-approved FBS and 100 μg/ml penicillin/streptomycin (Gibco) at 37 °C and 5% CO_2_. U2OS, PDNC4 (a gift from Dr. Ben Black^55^), HCT-116^SMC2-mAID-mCherry^ (a gift from Dr. Masatoshi Takagi^39^), HeLa^EGFP-AID-Sororin^ (a gift from Dr. Daniel Gerlich^56^) cells and mouse embryonic fibroblasts were cultured in DMEM high glucose (Gibco) supplemented with 10% Tet-approved FBS, 100 μg/ml penicillin/streptomycin, and GlutaMAX supplement (Gibco).

HCT-116^SMC2-mAID-mCherry/SMC3-TID^ cell line is derived from HCT-116^SMC2-mAID-mCherry^(39). Tagging of the endogenous locus of SMC3 was done according to the CRISPaint protocol^57^ using 2.5 μg frame selector plasmid (pCAS9-mCherry-Frame+2; Addgene_66941^57^), 2.5 μg target selector plasmid (pCS446_pSPgSMC3; this study) harboring SMC3 guide RNA (TGATACCACACATGGTTAATTGG), and 5 μg donor plasmid (pCS452_pCRISPaint-TurboID-pPGK-Hygro; this study). Cells were transfected with Fugene using standard procedures and subsequently selected using Hygromycin (Roche, 10843555001). U2OS cell line expressing H2B-mNeon/mCherry-CENP-A and HeLa^EGFP-AID-Sororin^ cell line expressing H2B-MNeon were obtained by lentiviral transduction (plasmids pLV-H2B-Neon-ires-Puro^58^ and pLenti6-CENP-A-mCherry^59^, Addgene_89767) using standard procedures and subsequently FACS-sorted (single cells) based on mCherry and/or H2B-mNeon expression.

Chicken DT40 (B lymphoma) cells were cultured in RPMI1640 medium supplemented with 10 % fetal bovine serum and 1% chicken serum at 39**°**C in 5% CO_2_ in air. Stable transfection of DT40 cells was performed as described previously^60^ to randomly integrate OsTIR1 into the genome of WTCDK1^as^ cell line^3^ to create WTCDK1^as^_TIR1 cell lines which was resistant to G418 1.5 mg/ml (Thermo fisher Scientific). TIR1 highly expressing clones were selected by western blotting using an antibody against TIR1 (a gift from Dr. Masato Kanemaki). In order to create SMC2-AID/CDK1^as^ or SMC3-AID/CDK1^as^ cell lines, we utilized Neon setting 24 (ThermoFisher Scientific). A plasmid encoding mAID-Clover with homologoues arms (2 µg) and a plasmid encoding hCas9 and Guide RNA (6 µg) were transfected into 2-4 million cells suspended in 100 µl buffer R from the Neon kit. Hygromycin 0.8 mg/ml (Thermofisher scientific) was used to select mAID-Clover tag integrated cell lines. Detailed protocols to establish DT40-AID/CDK1^as^ cell lines are available upon request to Dr. Kumiko Samejima.

### Treatments on human cells

Cells were arrested in G_2_ by addition of RO-3306 (10 μM; Tocris Bioscience; 4181) overnight and released in fresh media or in the presence of drugs, as indicated. Thymidine (2mM; Sigma-Aldrich; T1895) was added for 24 hours and cells were used for experiments between 6–10h after thymidine wash-out. In SMC2 and Sororin depletions, 3-indoleacetic acid (IAA) (500 μM, Sigma-Aldrich I5148-2G) was added overnight and present in the media in all the subsequent steps. 24 hours before IAA treatment, HCT-116^AID-mCherry-SMC2^ cells were supplemented with doxycycling (1 μg/ml; Sigma-Aldrich; D9891) to induce the expression of OsTIR1. In TurboID experiments, Biotin (500 μM,Sigma-Aldrich; B4501-5G) was added to the media 30 minutes before fixation. For knockdown experiments, 100 nM siRNA against RAD21 (SMARTpool: ON-TARGET plus Dharmacon L-006832-00-0010) or 100 nM of siGAPDH (Dharmacon, D-001830-01-05) were transfected in RPE-1 cells using RNAi Max (Thermo Fisher Scientific) according to manufacturer’s instructions. After 72 h of siRNA treatment, cells were cold-treated for 5 min, pre-extracted and fixed. Other treatments: Nocodazole (3.3 μM or 330 nM; Sigma-Aldrich M1404); MG-132 (5 mM; C2211).

### Plasmid construction

The list of primers used in this study can be found in Supplementary Table 1. The gRNAs for SMC3 was cloned into the vector pSPgRNA (Addgene_47108^61^) using primers H567/H568, as described previously^62^. pCS452_pCRISPaint-TurboID-pPGK-Hygro was created by Gibson cloning in two steps. First, TurboID contruct derived from 3xHA-TurboID-NLS_pCDNA3 (Addgene_107171^38^) was cloned into the BamHI site of plasmid CRISPaint-mNeon (addgene_174092^57^) with primers H574/H575 to create pCS451_CRISPaint-TurboID-mNeon. Next, mNeon was substituted with pPGK-Hygro derived from pMK290 (Addgene_72828^63^) using primers H582-H585.

### Immunofluorescence

For immunofluorescence, cells grown on 12 mm coverslip (no. 1.5H) were permeabilized for 1 min with warm 0.5% triton in PHEM buffer, followed by fixation for 10 min with 4% PFA (Electron microscopy sciences, 15710) in PBS. For analysis of cold-stable microtubules, cells were placed for 5 min on ice water prior to pre-extraction and fixation. After fixation, coverslips were washed three times with PBS and blocked with 3% BSA in PBS for 1h at room temperature. Primary antibodies diluted at 1:100 in 3% BSA were added to the coverslips and incubated for 4 hours at RT (a list with the primary and secondary antibodies can be found in Supplementary Table 1). Subsequently, cells were washed three times with 0.5 % triton in PBS and incubated with DAPI and secondary antibodies diluted 1:100 in 3% BSA for another 2 hour at RT. Next, coverslips were washed three times with 0.1% Triton and prepared for ExM or mounted onto glass slides using Prolong Gold antifade.

### Expansion Microscopy (ExM)

For ExM^23^, stained samples were treated with 0.1 mg/ml Acryloyl-X (Thermo Fisher A20770) in PBS for 2 hours, washed three times with PBS and incubated for 5 min in monomer solution (1×PBS, 2M NaCl, 2.5% (wt/wt) acrylamide 0.15% (wt/wt) N,N’-methylenebisacrylamide 8.625% (wt/wt) sodium acrylate). Coverslips were placed on top of a drop of 90 μl of freshly prepared gelation solution (monomer solution supplemented with 0.2% (wt/wt) TEMED and 0.2% (wt/wt) APS) and incubated for 1 hour at 37 °C. Gels were then incubated in digestion solution (8 units/ml proteinase K, 1×TAE, 0.5% TX-100, 0.8M guanidine HCl) for 2 hours at 37 °C and washed in PBS containing DAPI. For partial ExM (Fig. 4b), gel was washed several times with PBS (∼2-fold expansion). Full expansion was performed in a 10-cm plate by several washings of 30 minutes with excess volume of Milli-Q water (∼4.5-5-fold expansion). Expanded samples were immobilized on 25 mm (No. 1.5H) coverslips covered with 0.01 % (w/v) poly-L-lysin (Sigma-Aldrich, P8920) and imaged.

### Microscopy and image processing

Images from Figures 1f,g, 2a,f, S2, 4a, S5a, and 5e were acquired on a deconvolution System (DeltaVision Elite Applied Precision/GE Healthcare) with a ×100/1.40-NA objective (Olympus) using SoftWorx 6.0 software (Applied Precision/GE Healthcare). Images were acquired as z-stacks at 0.2 μm intervals and deconvolved using SoftWoRx using SoftWoRx. Images from figures 1a, h, i, S1b,i, 5d and S6c were acquired on a a Nikon Ti-E motorized microscope equipped with a Zyla 4.2Mpx sCMOS camera (Andor) and 100×1.35-NA objective lens (Nikon). Images were acquired as z-stacks at 0.2 μm intervals. Images from Figures 2d and 4b,e were acquired on a Zeiss LSM900 Airyscan2 with a 63×1.3-NA and using the Multiplex SR-2Y mode. For each experiment, optimized sectioning calculated by Zeiss software was used. Images from figures S1g, S3, 5a and S6a were acquired on a Nikon Ti2-E Microscope equipped with a VT-iSIM Super Resolution unit (VisiTech), a Prime BSI Express sCMOS camera (Photometrix) and a TIRF 100×1.49-NA objective lens (Nikon). Images were acquired as z-stacks at 0.5 μm intervals in live-cell imaging and 0.1 μm intervals in fixed samples. Images from Nikon and Zeiss systems were deconvolved with Huygens Professional (v20.10) using up to 40 iterations of the Classic Maximum Likelihood Estimation (CLME) algorithm with theoretical PSF. 3D reconstructions were created with Imaris Software (v9.7.2) (Bitplane). Confocal and Stimulated Emission Depletion Microscopy (STED) images were taken with an abberior Instruments STEDYCON STED setup equipped with an inverted IX83 microscope (Olympus), an APO100×1.4-NA oil objective and a pulsed 775 nm STED depletion laser. Star Orange was imaged with a pixel size of 25 nm and a pixel dwell time of 10 µs, using STED laser power of 100% and 640-nm excitation laser power of 29%. All acquisition operations were controlled by the STEDYCON Software (abberior Instruments).

### Live-cell imaging

For live-cell imaging, cells were seeded in 24-well glass bottom plates (No. 1.5H) (Cellvis, P24-1.5H-N). High-resolution live-cell imaging of U2OS^mCherry-CENP-A/H2B-mNeon^ cells was performed over 16 z-slices separated by 500 nm every 10 seconds on a Nikon Ti2-E Microscope equipped with a VT-iSIM Super Resolution unit (see above). Images were deconvolved with Huygens Professional (v20.10) using the conservative profile.

Imaging of HeLa^H2B-mNeon^ cells was performed over 5-8 z-slices separated by 2 mm every 4 min on an Andor CSU-W1 spinning disk (50 µm disk) with 30× 1.05-NA sil objective (Olympus) and an additional ×1.5 lens in front of an EMCCD camera. In both microscopes, cells were kept at 37 °C and 5% CO_2_ using a cage incubator and Boldline temperature/CO2 controller (OKO-Lab).

### Stretched chromosomes

Cells were seeded in 6-well plates at 50% confluency and incubated in 2 mM thymidine for 24 hours and released in 3.3 μM nocodazole overnight. In the morning, cells were harvested by mitotic shake-off, washed with PBS and resuspended in 15 ml of pre-warmed 0.4% Citrate solution. Cells were then incubated at 37°C for 20 mins and a 500 μl aliquot was cytospun (1200 rpm for 10 min at high acceleration) onto coated slides (Thermoscientific; 5991055). Cells were fixed in 4% PFA and immunostained.

### Image analysis

ImageJ (https://imagej.nih.gov/ij/)/Fiji (http://fiji.sc/<u>#</u>) (National Institute of Health) plot profile tool was used to measure intensity profiles, and intensities were normalized between 0 and 1. Quantification of number of centromere subdomains was manually scored in z-stacks using Fiji. Distance between CENP-A subdomains was measured with Imaris Software (v9.7.2) (Bitplane). Individual subdomains were detected using the spot detection tool set at an estimated XY diameter of 0.150 μm for ExM images and 0.100 μm for iSIM images. Next, distance between center of spots was measured with the measurement point tool. To minimize the error derived from axial chromatic aberration, only centromeres with subdomains that shared the same approximate focal plane were used for quantification. Images presented in the figures were only adjusted in brightness and contrast on raw or deconvoluted data using the Fiji software.

### Statistics and reproducibility

Statistical analysis was performed with Prism 9 Software (Graphpad software). Source data can be found in Supplementary Table 2. Representative results are displayed or all data were reported, as specified in individual figure legends. Data are presented as mean ± s.d. The comparisons most pertinent for the conclusions and number of independent experiments are specified in the figures and legends. Source data is provided in Supplementary Table 2. Data are presented as mean ± s.d. All replicates showed similar results and a representative experiment was reported.

### CENP-A ChIP-seq experiments and analysis

Nuclei were isolated from 1.5 × 10^9^ DT40 cells and digested with 60 units/ml MNase (Takara Bio Inc.) in buffer A (15 mM Hepes-KOH, pH 7.4, 15 mM NaCl, 60 mM KCl, 1 mM CaCl2, 0.34 M sucrose, 0.5 mM spermidine, 0.15 mM spermine, 1 mM DTT, and 1× complete protease inhibitor cocktail; Roche). After centrifugation at 17,800 g for 5 min, the chromatin pellet was suspended with buffer B (20 mM Tris-HCl, pH 8.0, 0.5 M NaCl, 10 mM EDTA, and 1× complete protease inhibitor cocktail; Roche), and then mononucleosome was extracted. The extracted mononucleosome fraction was incubated for 2 h at 4°C with Protein G Sepharose beads (GE Healthcare), which were preincubated with rabbit polyclonal anti–chicken CENP-A antibody^64^. Beads were washed with buffer B four times, and the bound DNA was purified by phenol-chloroform extraction and ethanol precipitation. The purified DNA was analyzed on a DNA sequencer (HiSeq 2500; Illumina). ChIP-seq libraries were constructed with the TruSeq DNA LT Sample Prep kit (Illumina) as described in the protocols provided with the kit. In brief, ∼50 ng of purified DNA was end-repaired, followed by the addition of a single adenosine nucleotide at 3’ and ligation to the universal library adapters. DNA was amplified by eight PCR cycles, and the DNA libraries were prepared. ChIP DNA libraries were sequenced using the HiSeq 2500 in up to 2 × 151 cycles. Image analysis and base calling were performed with the standard pipeline version RTA1.17.21.3 (Illumina).

Single-end reads (100bp) were trimmed with Trimmomatic v0.36 to remove adapters and low-quality bases. Trimmed reads were aligned to the galGal6 reference genome using BWA mem v0.7.16 and unique alignments were selected for downstream analysis. Read coverage profiles were generated with deepTools bamCoverage v3.5 using bins per million mapped reads (BPM) normalisation.

### Capture-C experiment and analysis

Careful analysis in DT40 cells reveals that the size of the core centromere remains mostly constant but the centromere position can drift over time in culture. (Importantly, it does not shift for any significant distance in cultures grown for <2 weeks grown from a single cell). Therefore, we performed CENP-A ChIP-seq to define the position of core centromeres in each subclone of wt/CDK1^as^, SMC2-AID/CDK1^as^ and SMC3-AID/CDK1^as^ cell lines. The Zcen position in the wt and SMC2-AID subclones was similar (42,767-42,792 kb and 42,767-42,797 kb respectively, based on the Galgal6 genome annotation. (Fig. S5c). In our SMC3-AID/CDK1as subclone the centromere position had shifted towards the q arm side by ∼10 kb (42,777-42,803 kb). Based on this mapping, we designed 34 viewpoints spaced 5-6 kb apart, covering 217 kb surrounding the ∼30 kb CENP-A region at the core of the Zcen (Supplementary Table 3). Each viewpoint corresponds to a DpnII-digested fragment with 2 capture-oligos complementary to either end of the fragment. To avoid the effect of non-cleaved restriction sites, 1-3 kb surrounding each fragment cannot be used for the interaction map. Therefore, the oligos were split into 3 pools (A, B, C) to space the fragments. further apart. In order to follow structural change of centromeric chromatin upon mitotic entry, wt CDK1as cells were treated with 1NMPP1 (13 h) to arrest cultures in late G_2_ and cross-linked with 1% Formaldehyde. To look at cells in late-prometaphase, cells were also crosslinked 30 min after 1NMPP1 washout. In order to determine the effects of the mitotic spindle on kinetochores, 0.5 µg/ml nocodazole was added after 1NMPP1 washout and crosslinked after 30 min (sample = Mnoc). Cells were cross-linked with 1% formaldehyde in culture media, glycine was added and then washed with PBS. Cells were lysed with detergents and then DNA was digested with DpnII, ligated, sonicated, and then made into a 3C library. This 3C library was subjected to pull-down twice with each capture-oligo pool to enrich the fragments of interest. 150 bp of pair-wise sequence was obtained from the ends of each fragment and analysed with the capC-MAP software^65^. 2-3 biological replicates (starting cell pellets are different) exist for all the data sets.

### Polymer physics modelling

For the polymer physics modelling, coarse-grained molecular dynamics simulations were performed, in which collections of molecules are represented by beads, which interact with phenomenological force fields and move according to Newton’s laws^66^. More specifically, chromatin fibres and chromosomes were modelled as bead-and-spring polymers, while bridging proteins representing, for instance, condensin or cohesin complexes were represented by additional individual beads. We used the multi-purpose molecular dynamics package LAMMPS (Large-scale Atomic/Molecular Massively Parallel Simulator^67^). In this section we detail the potentials underlying the force fields used in the simulations.

### The chromatin fibre

A chromatin fibre, corresponding to a 440 kbp chromatin region surrounding the core centromere, was discretised as a set of monomers, each of size corresponding to 1 kbp, or to a physical size σ∼ 10-20 nm, which could in principle be determined by fitting to microscopy experiments. There were three types of chromatin beads: one representing pericentromeric chromatin (PC), one representing the core centromere (CM) and one representing loop bases (LB) in PC. Below we specify when the potentials used depend on chromatin bead type. When nothing is specified, the potential is applied regardless of bead type.

Any two monomers (*i* and *j*) in the chromatin fibre interact purely repulsively, via a Weeks-Chandler-Anderson (WCA) potential, given by

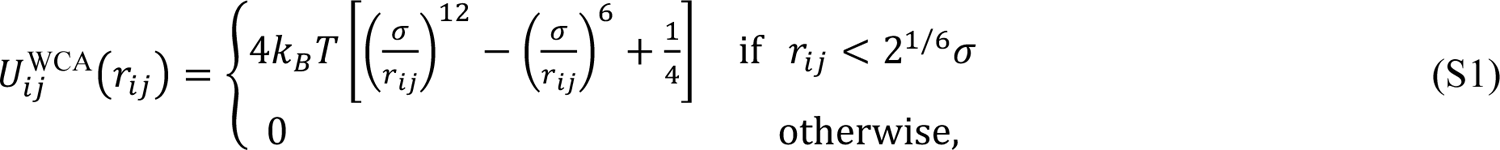

where *r_ij_* is the separation of beads *i* and *j*. There is also a harmonic elastic spring acting between consecutive beads in the chain to enforce chain connectivity. This is given by

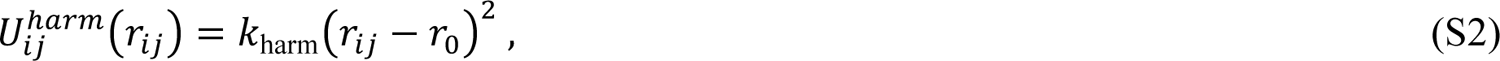

where *i* and *j* are neighbouring beads, *r*_0_ = 1.1σ is the maximum separation between the beads, and *k*_harm_ = 100 *k*_B_T⁄σ^2^ is the spring constant. Additionally, a triplet of neighbouring beads in the chromatin fibre interact via a Kratky-Porod term to model the stiffness of the chromatin fibre, which explicitly reads as follows,

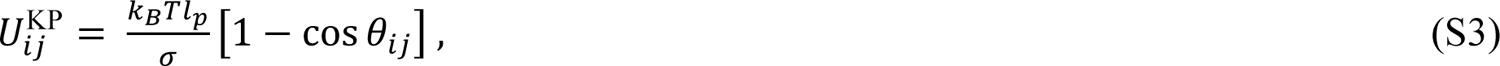

where *i* and *j* are neighbouring beads, while θ_ij_ denotes the angle between the vector connecting beads *i* and *j = i+1* and the vector connecting beads *j* and *j+1*. The quantity *l*_p_ is related to the persistent length of the chain. We set it to 3σ in our simulation for adjacent beads, which corresponds to a relatively flexible chain. For model 3, we set this value to 10σ for beads in the CM, which means that the latter region is locally stiffer in those simulations.

### Chromatin bridges

An important component of the model is constituted by multivalent chromatin-binding proteins, or bridges, which can bind to chromatin at multiple places. Chromatin bridges were modelled as spheres, again with size σ for simplicity. While these may represent a number of factors, we identified the two bridges simulated with condensin and cohesin (see main text). This is motivated by recent work showing that SMC proteins have a bridging activity alongside a chromatin looping (or extrusion) activity^34^.

In our simulations, the interaction between a chromatin bead, *a*, and a multivalent chromatin bridge, *b*, was modelled via a truncated and shifted Lennard-Jones potential, given by

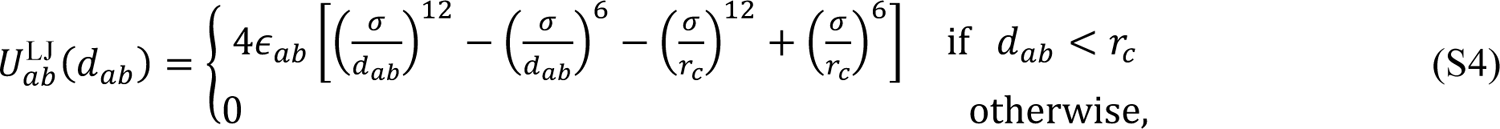

where *d*_ab_ denotes the distance between the centres of the chromatin bead and protein, *r*_c_ = 1.8 σ is a cut-off parameter, and ∈_ab_ determines the interaction strength between the bridge and chromatin. Note this is not equal to the value of the potential at the minimum point in view of the shift. The values of ∈_ab_ chosen are discussed below (see *Parameter values*). When bridges did not stick to a given chromatin region, the corresponding potential was instead set to a WCA potential as in Eq. (S1), corresponding to only steric interactions. Bridges also interacted with each other via steric interaction only (WCA potential).

Bridges could switch back and forward between a binding and a non-binding state with a rate *k*_sw_. This feature mimics post-translational modifications on protein complexes and accounts for the dynamical turnover of constituents within nuclear protein clusters^35^. When bridges were non-binding, they interacted with chromatin beads via the WCA potential.

### Parameter values and details of specific models

While general parameter values used in all simulations are mentioned above, here we list the model-specific parameters we used. The values of ∈_ab_ chosen for each case as given in Supplementary Table 4 (in units of *k*_B_T). In Model 0, we considered a single bridge (representing condensin, or Multivalent Protein 1 (MP1) in the main text), which could bind to PC more weakly than to the CM. This resulted in an “inverted” Capture-C signal with respect to our mitotic one, and to a compact configuration with the CM internal. In Model 1, we consider two bridges, with condensin binding more strongly to PC than to the CM. This led to a qualitatively correct Capture C signal, and to a switch in the CM location, which was now peripheral. In Model 0 and 1, no switching was considered. In Model 2, we used parameters as in Model 1, but both bridges could switch at a rate *k*_sw_ = 0.001 τ_B_^-1^, where the Brownian time τ_B_ = σ^2^/*D*, with *D* the diffusion coefficient of the beads. The Brownian time gives an order-of-estimate measure of the time it takes for a bead to diffuse across its own diameter, and it is this timescale which one can use to determine the mapping of simulation time to real time where required.

In Model 3, we also included chromatin loops, mimicking the looping activity of SMC proteins (and arising for instance due to loop extrusion and important for mitotic chromatin formation^3,36^). We included for simplicity fixed chromatin loops. Chromatin loops in a simulation were distributed randomly in PC, according to a Poisson distribution with average loop size equal to 40 kbp. We simulated over a number of runs (typically 50 for each condition), and each run was performed with a different set of random loops. For these runs, the two loop bases corresponding to each chromatin loop interacted with a harmonic spring, Eq. (S2), with *r*_0_ = 1.8σ and *k*_harm_ = 100 *k*_B_T⁄σ^2^. These loops create a bottlebrush topology. To control the stiffness of the backbone of the bottlebrush (consisting in the bases of consecutive loops), we included a Kratky-Porod potential, Eq. (S3), between consecutive LB beads, with *l*_p_ = 10σ. Regarding bridges, when modelling mitotic conditions, we considered two cases. In the first, we only included condensin-like bridges, which could bind strongly to LBs in PC, moderately to other PC beads, and weakly to the CM (Fig. 3f,g of the main text). We also performed additional simulations including cohesin (or MP2 in the main text), modelled as an additional bridge with moderate interactions to PC and stronger affinity for the CM. To represent G_2_ conditions, we modelled cohesin activity as a strong interaction with the CM and for simplicity we neglected interactions between condensin and CM and between cohesin and PC. In model 3, we considered switching bridges with rate *k*_sw_ = 0.001 τ_B_^-1^. The only exception was the G_2_ condition, where the cohesin-like bridge did not switch.

Numbers of bridges were chosen as follows: 100 (condensin) in model 0, 100 (condensin) in model 1, 200 (condensin) in model 2, 100+100 (condensin+cohesin) in model 3. When switching was included, half of the bridges of any type were active on average at any given time.

In model 3, we found a transition, or crossover, between a mitotic-like state (M) and an interphase-like state (G_2_). The former has typically a bipartite structure with condensin-rich clusters separated by a largely unstructured CM. In the latter, cohesin forms an additional cluster at the CM. The transition between the M and the G_2_-state can be triggered, for instance, by increasing the affinity between the cohesin-like bridge and the CM.

### Additional simulation details

To simulate Capture-C directionality curves, we identified contacts made by each chromatin bead (say bead *i*) with all other beads *j*. Two beads were considered to be in contact if their 3D distance was below a cross-linking threshold, which was varied for each condition to fit the experimental data (with the exception of model 0, which is not qualitatively right so that fitting is not possible). From contact data, directionality curves were drawn by computing for each bead *i* the natural logarithm of the ratio between the interactions in the region to the left of *i* and the interactions to the right of *i*. For both right and left interactions, contacts were summed up between a lower and an upper end threshold, equal to 3 kbp and 100 kbp respectively. Thresholds used for each model/condition were as follows: 5σ (model 0, 1, 2); 16σ (model 3; mitosis); 9σ (model 3; G_2_); 12σ (model 3; mitosis with additional cohesin). Pearson correlations between experimental and simulated Capture-C directionality curves were as follows: −0.77 (model 0); 0.95 (model 1); 0.97 (model 2); 0.97 (model 3, mitosis); 0.69 (model 3, G_2_; 0.96 in the region between −50 and 50 kbp from the middle of the CM); 0.98 (model 3, mitosis with additional cohesin).

To find the fraction of bipartite and monopartite structures, we analysed configurations from all simulations. The clustering algorithm in (^68^) was used to find clusters of condensin bridges (or active condensin bridges in models with switching), with a threshold of 3σ for the 3D distance of any two bridges to be in the same cluster. Error bars were determined by assuming Poisson distributions.

In all simulations, the core CM was modelled as an 11 kbp region in the middle of the simulated polymer. As epigenetic marks and histone binding is not solely determined by sequence, we assumed CENP-A (orange in the snapshots) to bind an additional 10 kbp region on either side in the visualisation – in this way the CENP-A-covered region corresponds approximately to the centromere size which is defined biologically (see gray region in the experimental Capture-C plots).

## Acknowledgements

We thank Masatoshi Takagi, Daniel Gerlich, Ben Black, Masato Kanemaki and Don Cleveland for reagents. We thank Abberior Instruments for imaging on STEDYCON system. The Hubrecht Flow Cytometry facility for help with sorting and the Hubrecht Imaging Centre for help with microscopy. We thank the Kops and Earnshaw lab members for discussions and comments on the manuscript. Figures 1j, 2c,e,h, S3a, 4c and 5f were created with Biorender.com. This study was funded by the European Research Council (ERC-SyG 855158) and by the Netherlands Organisation for Scientific Research (NWO/OCENW.KLEIN.182). The Kops lab is part of the Oncode Institute, which is partly funded by the Dutch Cancer Society (KWF Kankerbestrijding). The Earnshaw lab is funded by a Wellcome Principal Research Fellowship to WCE (107022). The Wellcome Centre for Cell Biology is funded by Wellcome grant 203149. The Marenduzzo lab is funded by the European Research Council (ERC CoG 648050, THREEDCELLPHYSICS). The Gilbert lab is funded by the UK Medical Research Council (MR/J00913X/1; MC_UU_00007/13).

## Author contributions

C.S., K.S., W.C.E., N.G. and G.J.P.L.K. conceived the project. C.S., K.S., W.C.E. and G.J.P.L.K. wrote the manuscript. C.S. designed, performed and analysed microscopy experiments unless specified otherwise. K.S. and A.B designed and performed Capture-C experiments, which were analysed by D.R. using software written and interpreted by C.A.B.. L.A.R. and M.L.A.L. performed experiments in Fig. 1g and 2d, respectively. D.M. designed and performed modeling experiments. T.H. and T.F. performed ChIP-seq experiments which were analysed by S.W.

**Correspondence and requests for materials** should be addressed to Carlos Sacristan, Kumiko Samejima, William C. Earnshaw or Geert J. P. L. Kops.

**The authors declare no conflict of interests.**

## Notes

### Competing Interest Statement

The authors have declared no competing interest.

