## Supplementary Figures for "Condensin reorganizes centromeric chromatin during mitotic entry into a bipartite structure stabilized by cohesin"

Figure S1. Sacristan C., Samejima K., *et al.* 2022

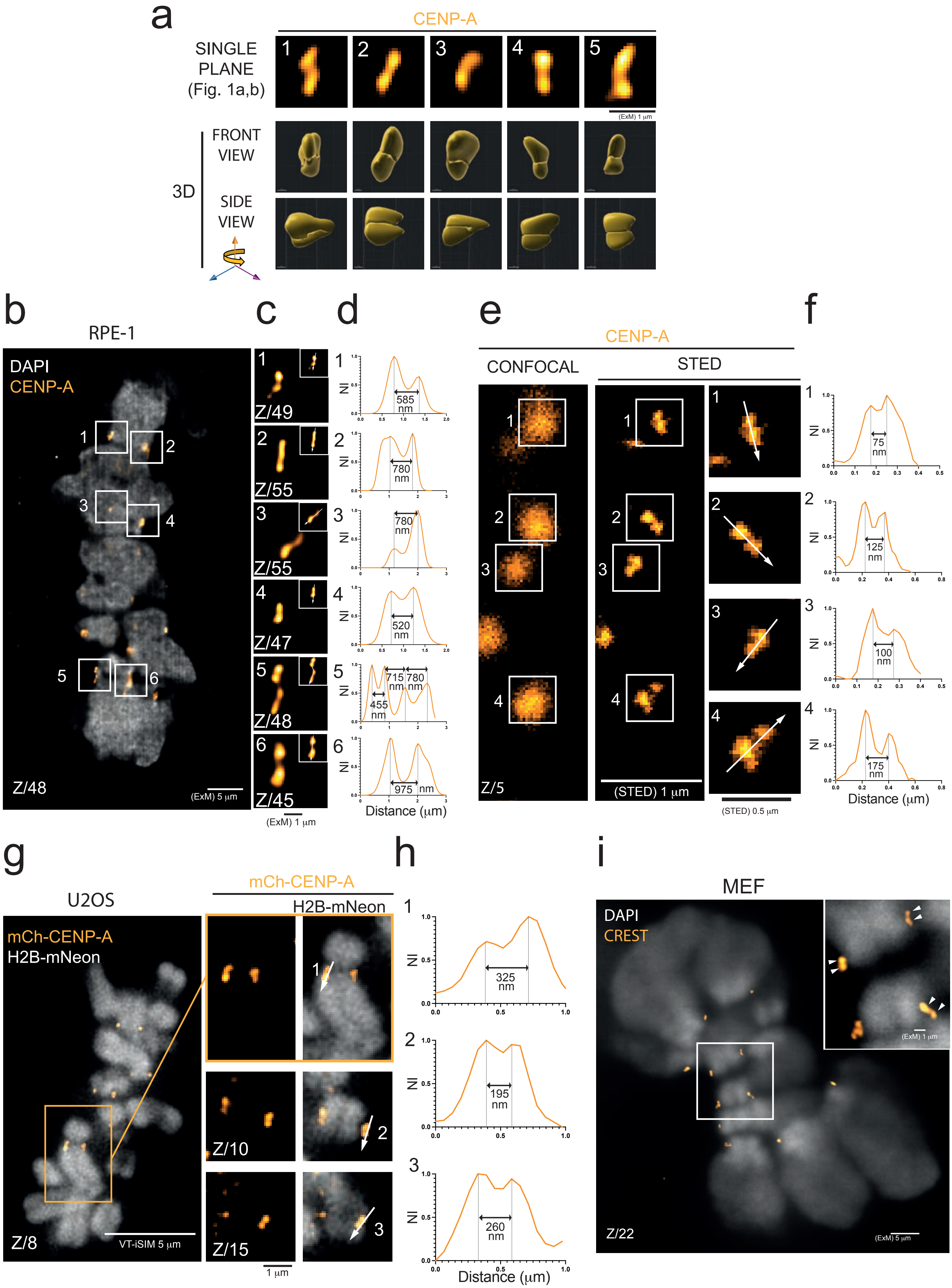

Figure S2. Sacristan C., Samejima K., *et al.* 2022

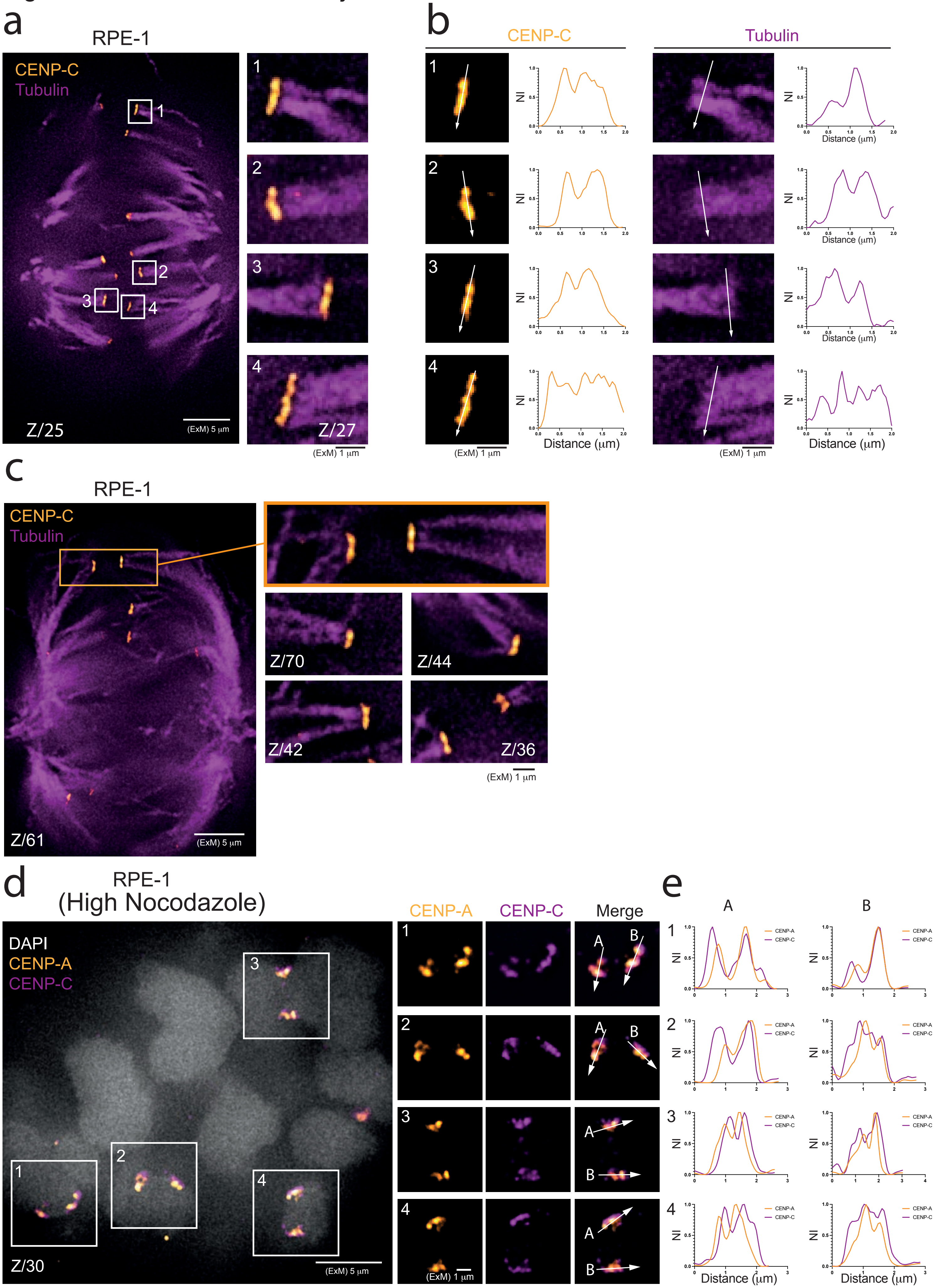

Figure S3. Sacristan C., Samejima K., *et al.* 2022

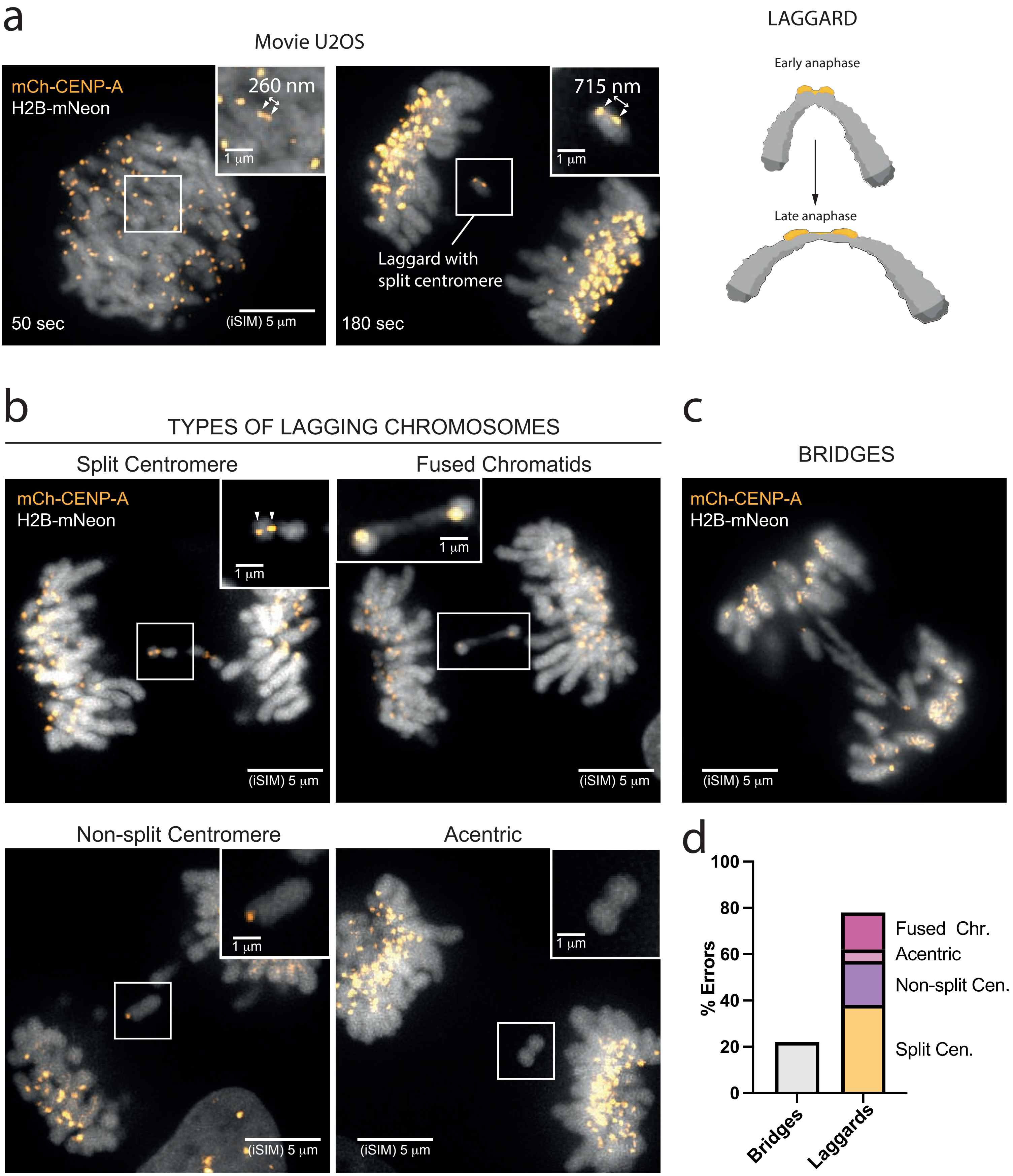

Figure S4. Sacristan C., Samejima K., *et al.* 2022

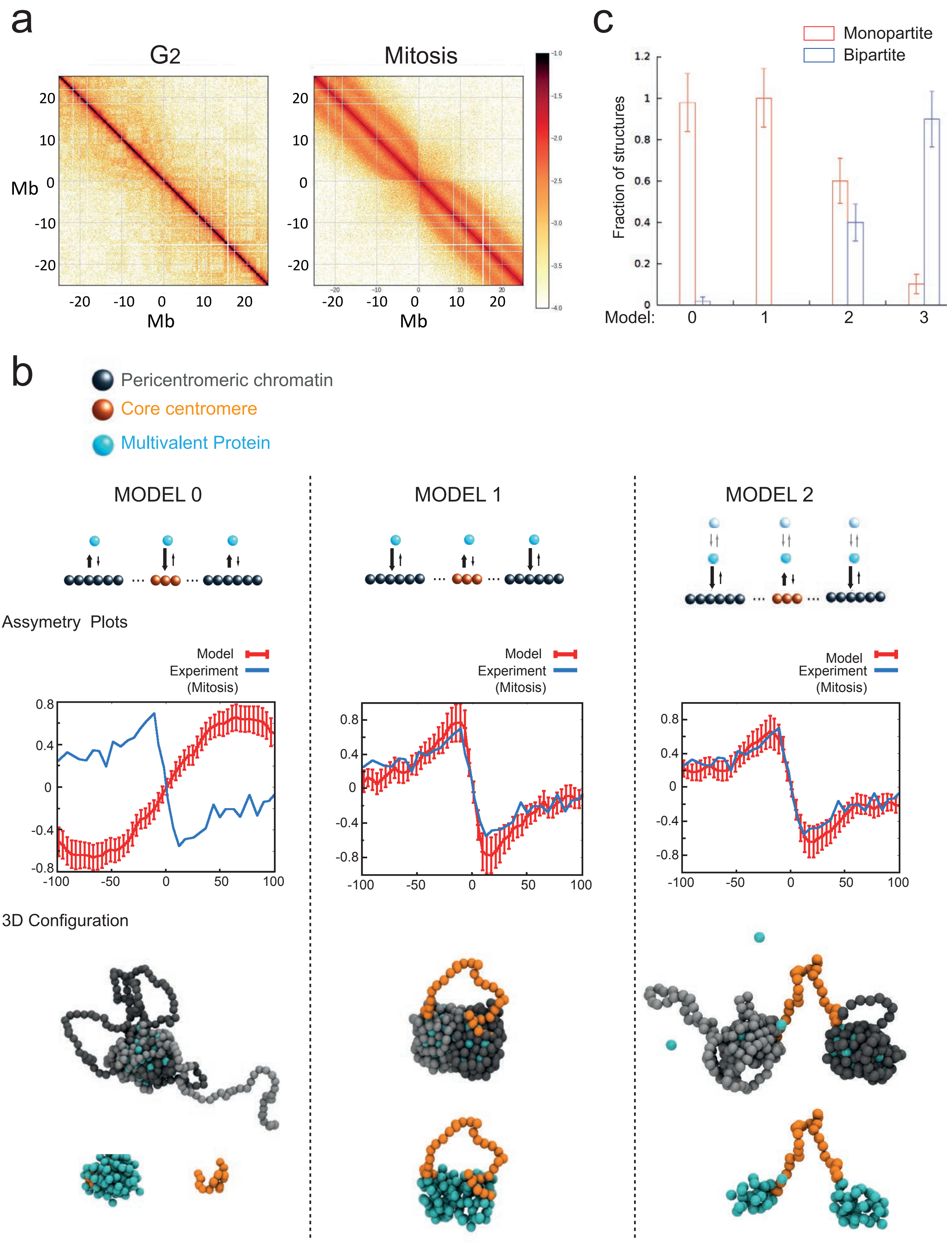

Figure S5. Sacristan C., Samejima K., *et al.* 2022

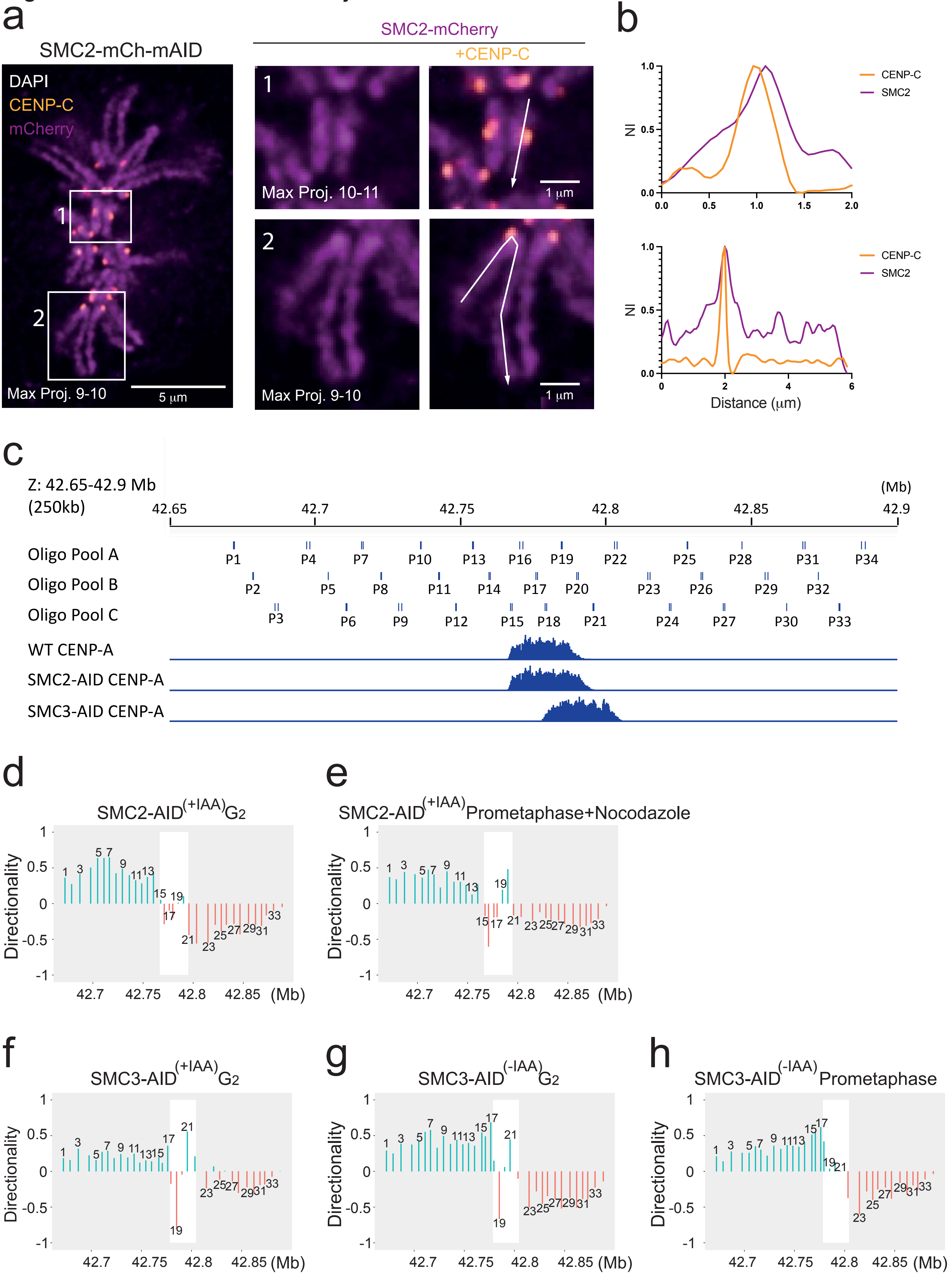

Figure S6. Sacristan C., Samejima K., *et al.* 2022

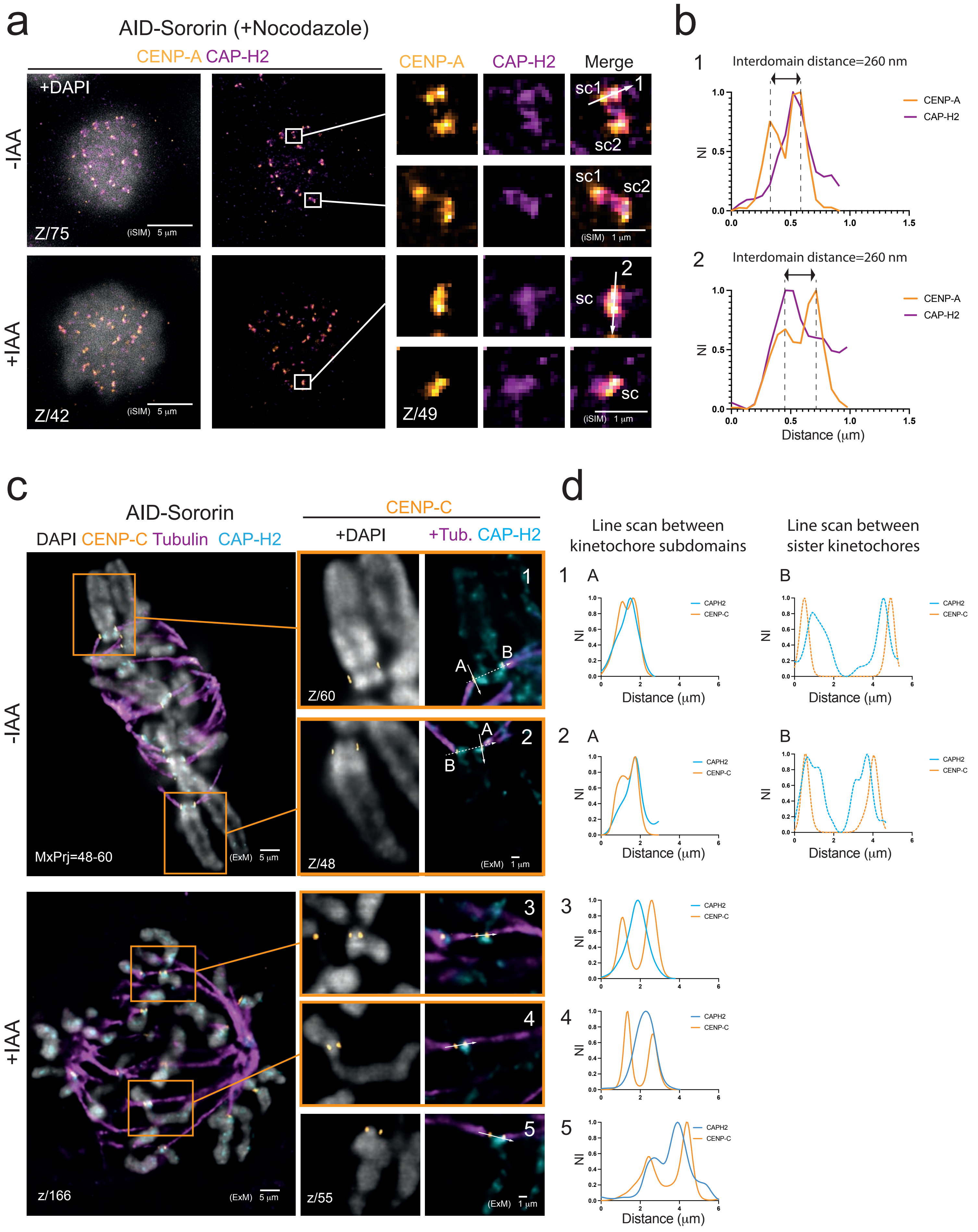
